# Identification of an endocannabinoid gut-brain vagal mechanism controlling food reward and energy homeostasis

**DOI:** 10.1101/2020.11.14.382291

**Authors:** Chloé Berland, Julien Castel, Romano Terrasi, Enrica Montalban, Ewout Foppen, Claire Martin, Giulio G. Muccioli, Serge Luquet, Giuseppe Gangarossa

## Abstract

The regulation of food intake, a *sine qua non* requirement for survival, thoroughly shapes feeding and energy balance by integrating both homeostatic and hedonic values of food. Unfortunately, the widespread access to palatable food has led to the development of feeding habits that are independent from metabolic needs. Among these, binge eating (BE) is characterized by uncontrolled voracious eating. While reward deficit seems to be a major contributor of BE, the physiological and molecular underpinnings of BE establishment remain elusive. Here, we combined a physiologically relevant BE mouse model with multiscale *in vivo* approaches to explore the functional connection between the gut-brain axis and the reward and homeostatic brain structures.

Our results show that BE elicits compensatory adaptations requiring the gut-to-brain axis which, through the vagus nerve, relies on the permissive actions of peripheral endocannabinoids (eCBs) signaling. Selective inhibition of peripheral CB1 receptors resulted in a vagus-dependent increased hypothalamic activity, modified metabolic efficiency, and dampened activity of mesolimbic dopamine circuit, altogether leading to the suppression of palatable eating. We provide compelling evidence for a yet unappreciated physiological integrative mechanism by which variations of peripheral eCBs control the activity of the vagus nerve, thereby in turn gating the additive responses of both homeostatic and hedonic brain circuits which govern homeostatic and reward-driven feeding.

In conclusion, we reveal that vagus-mediated eCBs/CB1R functions represent an interesting and innovative target to modulate energy balance and counteract food-reward disorders.

## Introduction

Feeding is a complex and highly conserved process whose orchestration results from the dynamic integration of interoceptive and exteroceptive signals. The homeostatic and hedonic components of feeding have been attributed to the hypothalamus and the dopamine (DA) reward system, respectively [1]. While the first can be broadly defined as the key regulator of food intake to ensure optimal energy balance, the second mainly relates to the reinforcing properties of sensory stimuli (perception, cues, taste, odors) and reward-associated features of feeding. However, despite the well-accepted recognition that both feeding components are tightly and functionally interconnected [1], they are usually investigated as isolated systems. In addition, the counterpointing central *vs* peripheral regulations of feeding add a supplemental degree of complexity in the identification of integrative regulatory mechanisms [2]. While energy homeostasis refers to negative feedback mechanisms maintaining body weight at *set-points*, the combination of both homeostatic and hedonic components of feeding leads to the establishment of feed-forward mechanisms of physiological adaptations. Feed-forward adaptation, also known as allostasis (*stability through changes*), is critical for energy balance and metabolic efficiency [3] but also contributes to reward-associated events [4]. Indeed, the widespread availability and consumption of palatable diets have profoundly altered the delicate allostatic integration of homeostatic and hedonic signals, leading to the development of metabolic disorders. This is particularly evident in food reward-driven dysfunctions such as binge eating (BE), where uncontrolled feeding perfectly recapitulates the efforts for an organism to adapt its homeostatic processes to the hedonic aspects of feeding. In fact, short- and/or long-term consumptions of energy-rich palatable diets remodel the DA reward system [5] and promote functional adaptations within the hypothalamus [6–8]. Beyond these two core processors of feeding, recent reports have mechanistically demonstrated that the gut-brain vagal axis, besides sensing interoceptive signals and influencing feeding and energy homeostasis [9–11], is also a major modulator of the reward system [12–14]. However, the physiological processes by which the gut-to-brain axis modulates reward feeding remain unclear. Emerging evidence strongly suggest that, besides a plethora of peripheral hormones (*i*.*e*. ghrelin, leptin, GLP-1, CCK), peripheral endocannabinoids (eCBs) may be fundamental players in the regulation of feeding and metabolic efficiency [15–17]. Indeed, eating disorders-associated alterations in peripheral eCBs have been reported in obese and BE patients [18, 19] as well as in diet-induced obese rodents [15, 20]. However, whether and how peripheral eCBs play a permissive role in guiding reward-based feeding behaviors and in buffering the allostatic regulation of energy balance is unexplored.

To tackle this question, we took advantage of a physiologically relevant BE-like mouse paradigm which, by promoting anticipatory and escalated consummatory food responses, triggers reward-driven behavioral, molecular and allostatic adaptations. Binge eating, which elicited DA-dependent molecular modifications in the reward-related structures [dorsal striatum (DS) and nucleus accumbens (NAc)], revealed an unappreciated integrative gut-to-brain orchestration requiring the modulatory actions of peripheral eCBs. In particular, we show that BE requires an orchestrated dialog between peripheral eCBs and both central hypothalamic and VTA structures through the gut-brain vagal axis, thus modulating both energy balance and reward-like events.

## Material and methods

See **Suppl. Material** for the detailed description of materials, methods and techniques. See **Suppl. Table 1** for detailed statistics and experimental sizes.

### Animals

All experiments were approved by the Animal Care Committee of the Université de Paris (CEB-25–2016) and carried out following the 2010/63/EU directive. 8-12 weeks old male and/or female C57Bl/6J mice (Janvier, Le Genest St Isle, France) were single-housed one week prior to any experimentation in a room maintained at 22 +/-1 °C, with light period from 7 am to 7 pm. Regular chow diet (3 438 kcal/kg, protein 19%, fat 5%, carbohydrates 55%, of total kcal, reference #U8959 version 63 Safe, Augy, France) and water were provided *ad libitum. Drd2*-Cre mice [Tg(Drd2-cre)ER44Gsat/Mmucd, Jackson laboratory] were used for *in vivo* fiber photometry Ca^2+^ imaging in the VTA. *Drd2*-eGFP mice [Tg(Drd2-eGFP)S118Gsat/Mmnc] were generated by GENSAT. See **Suppl. Material** for further details.

### Behaviors

#### Palatable binge eating-like paradigm

Intermittent access to a palatable mixture (Intralipid 20% w/v + sucrose 10% w/v) was provided 1-hour/day during 10-14 consecutive days at 10-11 am. During binge sessions chow pellets were not removed. Volume (ml) of consumed palatable mixture was measured at the end of the session.

#### Locomotor activity

Locomotor activities were measured using an infrared beam-based activity monitoring system (Phenomaster, TSE Systems GmbH, Germany).

#### Tail suspension

To record the activity of GCaMP6f-expressing VTA neurons, mice were suspended above the ground by their tails. Ca^2+^ imaging was performed before and during tail suspension.

#### Exploratory drive in a new environment

To record the activity of GCaMP6f-expressing VTA neurons, mice were put in a new environment (NE) consisting in a novel cage. Ca^2+^ imaging was performed before and immediately after changing the environment. See **Suppl. Material** for further details.

### Metabolic efficiency analysis

Mice were monitored for whole energy expenditure (EE), O_2_ consumption, CO_2_ production, respiratory exchange rate (RER=VCO_2_/VO_2_, V=volume), and locomotor activity using calorimetric cages (Labmaster, TSE Systems GmbH, Bad Homburg, Germany). Ratio of gases was determined through an indirect open circuit calorimeter. This system monitors O_2_ and CO_2_ at the inlet ports of a tide cage through which a known flow of air is ventilated (0.4 L/min) and regularly compared to a reference empty cage. O_2_ and CO_2_ were recorded every 15 min during the entire experiment. EE was calculated using the Weir equation for respiratory gas exchange measurements. Food intake was measured with sensitive sensors for automated online measurements. Mice were monitored for body weight and composition at the entry and exit of the experiment using an EchoMRI (Whole Body Composition Analyzers, EchoMRI, Houston, USA). Data analysis was performed on Excel XP using extracted raw values of VO_2_, VCO_2_ (ml/h), and EE (kcal/h).

### Brown adipose tissue and telemetry body temperature measurements

#### Infrared camera for BAT temperature

Heat production was visualized using a high-resolution infrared camera (FLIR E8; FLIR Systems, Portland, USA). To measure brown adipose tissue (BAT) temperature, images were captured before and after binge sessions. Infrared thermography was analyzed using the FLIR TOOLS software.

#### Telemetric body temperature

Anesthetized mice were implanted with a telemetric transmitter (HD-XG; Data Sciences International) to longitudinally measure fluctuations of core temperature. They received a daily injection of ketoprofen (Ketofen® 10%) for 3 days. During a 7-days recovery period, mice were monitored and had facilitated access to food. Data were collected using the Ponemah® software. The detection of transmitted signals was accomplished by a radio receiver (temperature and activity) and processed by a microcomputer system.

### Tissue preparation and immunofluorescence

Mice were anaesthetized with pentobarbital (500 mg/kg, i.p., Sanofi-Aventis, France) and transcardially perfused with 4°C PFA 4% for 5 minutes. Sections were processed as previously described [21]. Quantification of immunopositive cells was performed using the cell counter plugin of ImageJ taking a fixed threshold of fluorescence as standard reference. See **Suppl. Material** for further details and list of antibodies.

### Western blotting and quantitative RT-PCR

At the end of the binge session, the mouse head was cut and immersed in liquid nitrogen for 3 seconds. Brains and sampled structures were processed as previously described [21]. See **Suppl. Material** for further details, list of antibodies and primers.

### Drug treatments

The following compounds were used: insulin (0.5 U/kg, Novo Nordisk), CCK-8S (10 μg/kg, Tocris), liraglutide (100 μg/kg, gift from Novo Nordisk), exendin 4 (10 μg/kg, Tocris), leptin (twice/day for 2 days at 0.25 mg/kg, Tocris), AM251 (3 mg/kg, Tocris), AM6545 (10 mg/kg, Tocris), JD-5037 (3 mg/kg, MedChemExpress), SKF81297 (5 mg/kg, Tocris), haloperidol (0.25 and 0.5 mg/kg, Tocris), SCH23390 (0.1 mg/kg, Tocris), GBR12909 (10 mg/kg, Tocris), d-amphetamine sulphate (2 mg/kg, Tocris), JZL184 (8 mg/kg, Tocris). See **Suppl. Material** for further details.

### Subdiaphragmatic vagotomy

Prior to surgery and during 3 post-surgery days, animals were provided with *ad libitum* jelly food (DietGel Boost #72-04-5022, Clear H_2_O). Animals received Buprécare® (buprenorphine 0.3 mg) and Ketofen® (ketoprofen 100 mg) and were anaesthetized with isoflurane (3.5% induction, 1.5% maintenance). Body temperature was maintained at 37°C. Briefly, using a binocular microscope, the right and left vagus nerve branches were carefully isolated along the esophagus and sectioned in vagotomized (VGX) animals or left intact in sham animals. Mice recovered for at least 3 weeks before being used for experimental procedures. See **Suppl. Material** for further details.

### Quantification of eCBs by UHPLC-MS/MS

Lipids were extracted by liquid/liquid extraction in the presence of deuterated standards. They were then purified by SPE to obtain a fraction containing eCB and NAE that were analyzed by UHPLC-MS/MS using a Xevo TQ-S (ESI source; positive mode). For each analyte, the signal (AUC) of the relevant internal standard was used for normalization. See **Suppl. Material** for further details.

### Viral production

pAAV.Syn.Flex.GCaMP6f.WPRE.SV40 (titer ≥ 1×10^13^ vg/ml, working dilution 1:5) was a gift from Douglas Kim (Addgene viral prep #100833-AAV9; http://n2t.net/addgene:100833; RRID:Addgene_100833). pAAV5-hSyn-dLigh1.2 (titer ≥ 4×10^12^ vg/ml) was a gift from Lin Tian (Addgene viral prep #111068-AAV5; http://n2t.net/addgene:111068; RRID:Addgene_111068).

### Stereotaxic procedure

Mice were anaesthetized with isoflurane, administered with Buprécare® and Ketofen®, and placed on a stereotactic frame (Model 940, David Kopf Instruments). pAAV.Syn.Flex.GCaMP6f.WPRE.SV40 (0.3 μl) was unilaterally injected into the VTA (L=™0.5; AP=™3.4; V=™4.4, mm) of *Drd2*-Cre mice at a rate of 0.05 μl/min. pAAV5-hSyn-dLight1.2 (0.5 μl) was unilaterally injected into the NAc (L=™0.75; AP=1.18; V=™4.4, mm) of C57BL6/J male and female mice at a rate of 0.05 μl/min. The injection needle was carefully removed after waiting 5 minutes at the injection site.

### Fiber photometry and data analysis

A chronically implantable cannula (Doric Lenses, Québec, Canada) composed of a bare optical fiber (400 μm core, 0.48 N.A.) and a fiber ferrule was implanted 100 μm above the viral injection site (VTA or NAc). The fiber was fixed onto the skull using dental cement (Super-Bond C&B, Sun Medical). Real-time fluorescence signals emitted by GCaMP6f or DA biosensor dLight1.2 [22] were recorded and analyzed as previously described [21]. Fluorescence was collected using a single optical fiber for both delivery of excitation light streams and collection of emitted fluorescence. The fiber photometry setup used 2 light-emitting LEDs: 405 nm LED sinusoidally modulated at 330 Hz and a 465 nm LED sinusoidally modulated at 533 Hz merged in a FMC4 MiniCube (Doric Lenses) that combines the 2 wavelengths excitation light streams and separate them from the emission light. The MiniCube was connected to a Fiberoptic rotary joint connected to the cannula. A RZ5P lock-in digital processor controlled by the Synapse software (Tucker-Davis Technologies, TDT, USA), commanded the voltage signal sent to the emitting LEDs via the LED driver (Doric Lenses). Data are presented as z-score of ΔF/F. See **Suppl. Material** for further details.

### Statistics

All data are presented as mean ± SEM. Statistical tests were performed with Prism 7 (GraphPad Software, La Jolla, CA, USA). Detailed statistical analyses are listed in the **Suppl. Table 1**. Normality was assessed by the D’Agostino-Pearson test. Depending on the experimental design, data were analyzed using either Student t-test (paired or unpaired) with equal variances, One-way ANOVA or Two-way ANOVA. The significance threshold was automatically set at p<0.05. ANOVA analyses were followed by Bonferroni *post hoc* test for comparisons only when overall ANOVA revealed a significant difference (at least p<0.05).

## Results

### Time-locked access to palatable diet induces adaptation of nutrients partitioning and metabolic efficiency

Several paradigms of bingeing are used to model eating disorders [23]. However, the majority of these paradigms rely on (*i*) prior alterations of basal homeostasis (food or water restriction/deprivation, stress induction), (*ii*) dietary exposure to either high-sugar or high-fat foods, or (*iii*) absence of food choice during bingeing periods. We therefore adapted existing protocols to better study reward and homeostatic components of food intake during binge eating (BE). Since dietary mixtures of fat and sugar lead to enhanced food-reward properties [24], we designed a highly palatable diet to promote intense reward-driven feeding. Time-locked access to this palatable diet was sufficient to drive escalating binge-like consumption without restricting access to chow diet (**Fig. 1A**). In that regard, this model is preferentially driven by reward values over metabolic demands since animals are neither food nor water restricted.

**Fig. 1:**
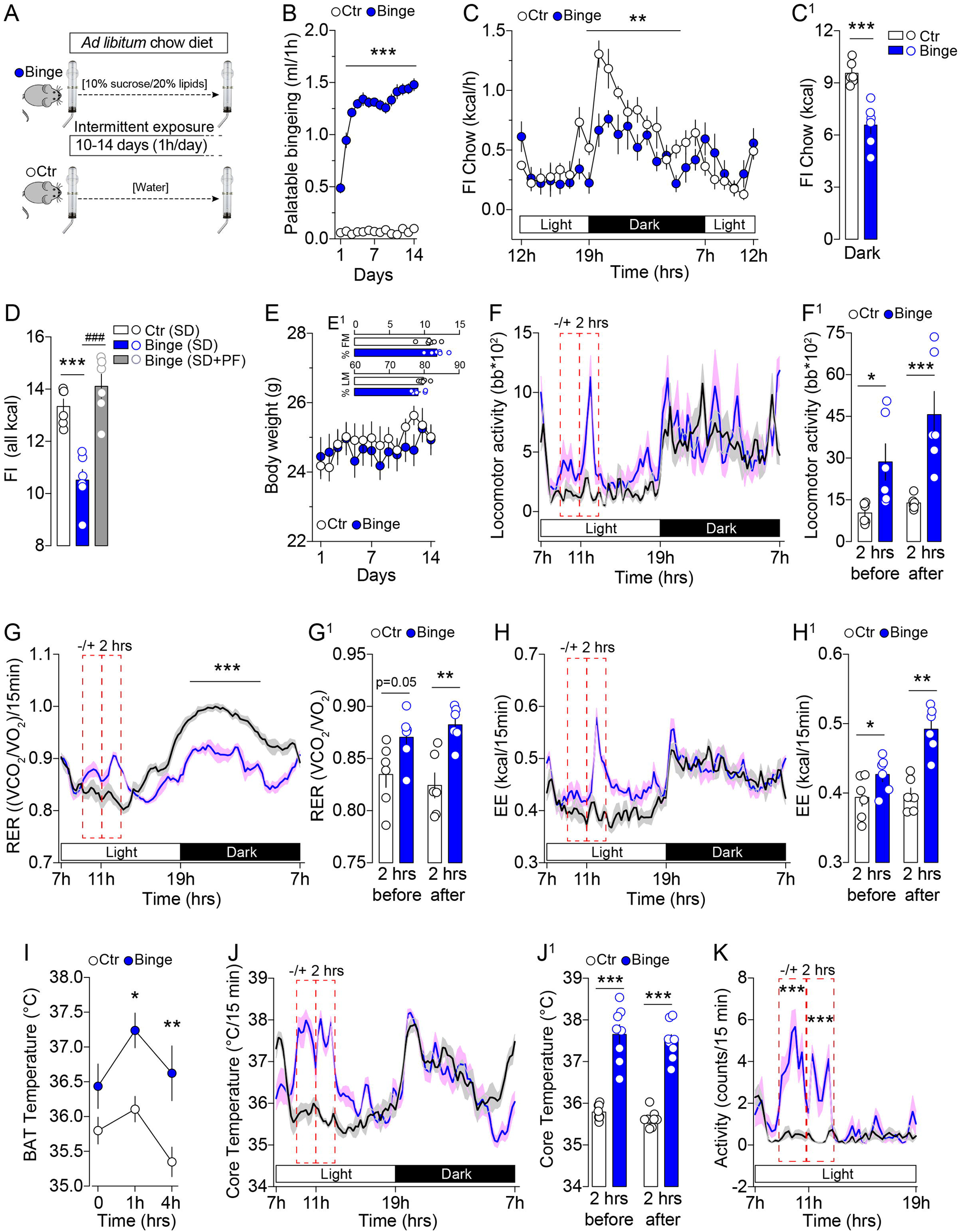
Allostatic adaptations of metabolic efficiency to binge eating. (**A**) Experimental design. Control (Ctr) or bingeing animals (Binge) had daily intermittent access to water or a palatable lipids/sucrose mixture for 1 hour/day during 10-14 consecutive days. Regular chow pellets and water were provided *ad libitum* throughout the entire experiment. (**B**) Daily binge consumption (ml) of palatable mixture during a 14-days protocol. Statistics: ***p<0.001 Binge *vs* Control. (**C**) Temporal pattern of regular chow food intake (FI, kcal/h) during 24 hrs (average of 3 consecutive days). Statistics: **p<0.01 Binge *vs* Control. (**C**^**1**^) Cumulative chow food intake during the dark period. Statistics: ***p<0.001 Binge *vs* Control. (**D**) 24 hrs food intake considering all calories: standard diet (SD) and palatable food (PF). Statistics: ***p<0.001 Binge(SD) *vs* Control(SD), ^###^p<0.001 Binge(SD+PF) *vs* Binge(SD). (**E**) Body weight throughout the 14-days experimental procedure. (**E**^**1**^) Body composition [% fat mass (FM) and % lean mass (LM)] of control and bingeing mice. (**F**) 24 hrs locomotor activity in calorimetric chambers (average of 3 consecutive days). Red dotted rectangles indicate the locomotor activity 2-hrs prior and after palatable food access. (**F**^**1**^) Cumulative locomotor activity 2-hrs prior and after palatable food access. Results are expressed as beam breaks (bb). Statistics: *p<0.05 and ***p<0.001 Binge *vs* Control. (**G**) Longitudinal profile of the respiratory energy ratio (RER) from indirect calorimetry (average of 3 consecutive days) and (**G**^**1**^) averaged RER values 2-hrs prior and after palatable food access. Statistics: **p<0.01 and ***p<0.001 Binge *vs* Control. (**H**) Longitudinal profile of energy expenditure (EE) from indirect calorimetry (average of 3 consecutive days) and (**H**^**1**^) averaged EE values 2-hrs prior and after palatable food access. Statistics: *p<0.05 and **p<0.01 Binge *vs* Control. (**I**) Brown adipose tissue (BAT) temperature during bingeing. Statistics: *p<0.05 and **p<0.01 Binge *vs* Control. (**J**) Real-time core temperature recording during 24 hrs and (**J**^**1**^) averaged values 2-hrs prior and after palatable food access. Statistics: ***p<0.001 Binge *vs* Control. (**K**) Matching locomotor activity from core temperature measurements. Statistics: ***p<0.001 Binge *vs* Control. For number of mice/group and statistical details see **Suppl. Table 1**.

Male and female mice intermittently exposed (bingeing) to this dietary palatable mixture maximized their intake within a few days (**Fig. 1B, Suppl. Fig. 1A**) and were characterized by a significant reduction in spontaneous chow food intake (**Fig. 1C, C**^**1**^, **Suppl. Fig. 1B**). However, the overall caloric intake [standard diet (SD) + palatable food (PF)] remained similar to controls, thus indicating a conserved maintenance in calories consumption despite reward-driven food intake (**Fig. 1D, Suppl. Fig. 1B**). Importantly, isocaloric feeding was associated with conserved body weight (BW) and body composition (**Fig. 1E, E^1^, Suppl. Fig. 1C**). Moreover, bingeing was simultaneously associated with an increased anticipatory locomotor activity ∼2 hours before food access and lasted for another ∼1-2 hours (**Fig. 1F, F^1^**), with no changes in the ambulatory activity during the dark phase (**Fig. 1F**).

Next, we investigated the consequences of BE on metabolic efficiency. Indirect calorimetry analysis revealed an increase in the respiratory exchange ratio (RER) before and after intermittent palatable food consumption (**Fig. 1G, G^1^**), whilst a stark reduction was detected in the dark phase (**Fig. 1G**), thereby highlighting a metabolic shift of energy substrates use (from carbohydrates to lipids as indicated by RER ∼1 or RER ∼0.7, respectively). This was further confirmed by the modulation of fatty acid oxidation (FAO, **Suppl. Fig. 1D**). In addition, we also observed an increased energy expenditure (EE) during the food anticipatory and consummatory phases (**Fig. 1H, H^1^**). Furthermore, infrared thermography revealed that BE was associated with a transient increase in brown adipose tissue (BAT) energy dissipation (**Fig. 1I**), while telemetric recording of core body temperature revealed a BE-specific increase during the anticipatory, consummatory and post-prandial phases (**Fig. 1J, J^1^**, **Suppl. Fig. 1E, E**^**1**^) and a sharp reduction during the last hours of the dark phase. Overall, changes in core body temperature were fostered around the time of locked palatable food access and overlapped with the increase in locomotor activity (**Fig. 1J, K**). Access to calories-rich food and time-restricted feeding are invariably associated with changes in circulating signals reflecting metabolic and behavioral adaptations [25]. In line with this, we observed that our BE model was associated with reduced circulating triglycerides and insulin, and increased circulating corticosterone during the anticipatory phase (**Suppl. Fig. 1F-H**), while glucose dynamics and glucose-evoked insulin release (oral glucose tolerance test) remained unchanged (**Suppl. Fig. 1I, J**). These data indicate that homeostatic adaptations occurring during time-locked palatable feeding lead to changes in lipid-substrates utilization and promote adaptive activation of the hypothalamic-pituitary-adrenal (HPA) axis. However, bingeing did not elicit major changes in the expression of key hunger- and satiety-related hypothalamic genes (*Npy, Agrp, Pomc, Cart, Hcrt*, **Suppl. Fig. 1K**). Overall, these results point to rapid reward-driven allostatic adaptations during which animals optimize their palatable food consumption and physiologically adapt to balance energy homeostasis and maintain a stable body weight.

### BE induces dopamine-related modifications in a D1R-dependent manner

Dopamine (DA)-neurons and DA-sensitive structures [dorsal striatum (DS) and nucleus accumbens (NAc)] are critical players in reward-based paradigms and BE disorders [26, 27]. Here, we investigated whether and how bingeing modulated the DA-associated signaling machinery. The phosphorylations of the ribosomal protein S6 and the extracellular signal-regulated kinases (ERK) were used as functional readouts of DA-dependent molecular activity [28, 29]. The food anticipatory phase was associated with an increase in P-ERK only in the DS (**Fig. 2A, B, Suppl. Fig. 2A-C**), mostly reflecting the increased locomotor activity during the anticipatory phase. Palatable food consumption induced an increase in P-ERK and P-S6 (Ser^235/236^ and Ser^240/244^ sites) in both DS and NAc (**Fig. 2A, B, Suppl. Fig. 2C**). Interestingly, acute (single) consumption of palatable diet failed to trigger P-ERK and P-S6 (**Fig. 2A, B, Suppl. Fig. 2C**), revealing that molecular adaptations of DA signaling in the DS/NAc require the full establishment of BE and not only palatable food consumption. Immunofluorescence analysis further confirmed BE-induced S6 activation (**Suppl. Fig. 2D, E**).

**Fig. 2:**
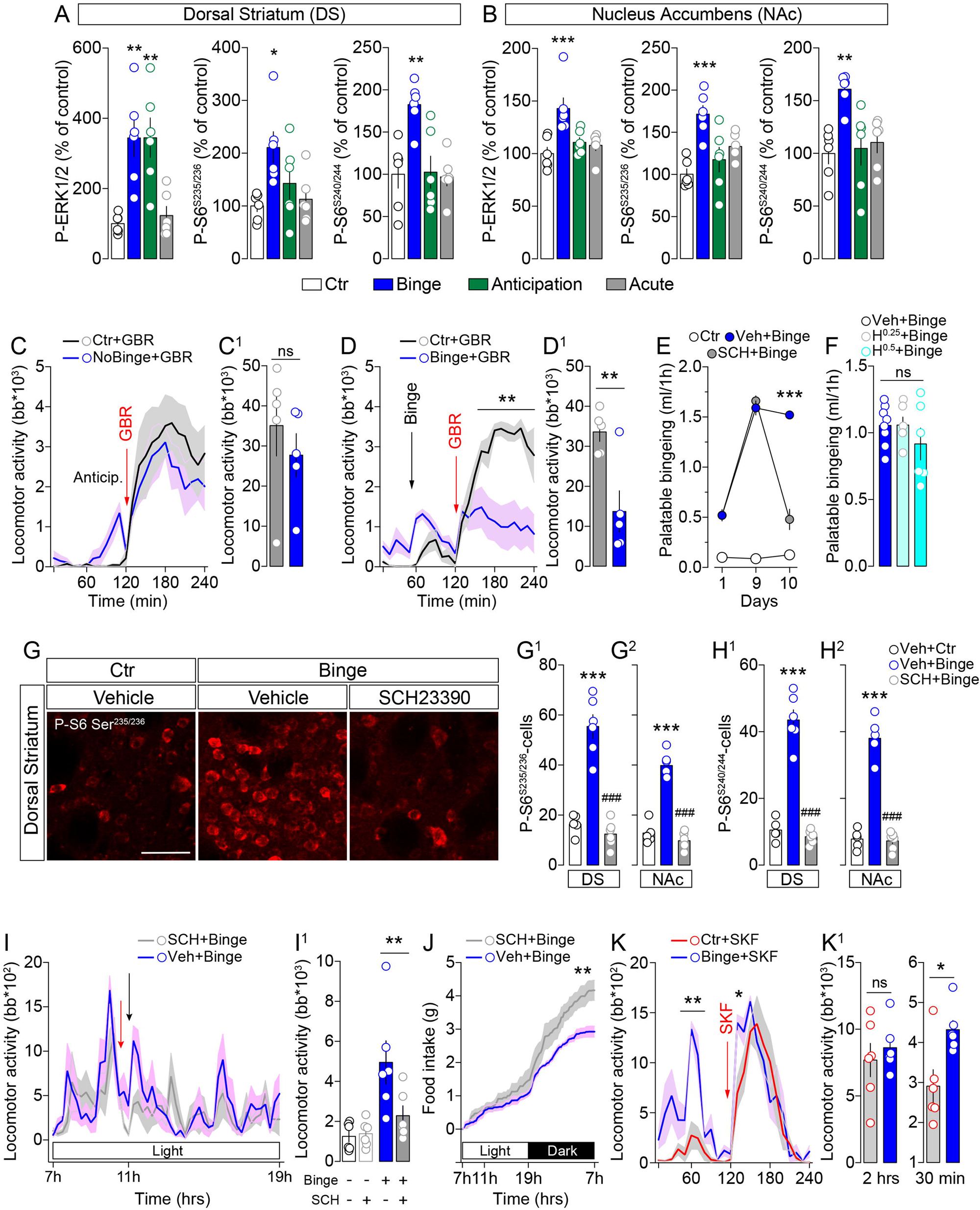
Binge eating induces dopamine D1R-related molecular modifications. (**A, B**) Protein quantification of phospho-ERK, S6^S235/236^ and S6^S240/244^ in the DS (**A**) and NAc (**B**). For immunoblot pictures see **Suppl. Fig. 2C**. Statistics: *p<0.05, **p<0.01 and ***p<0.001, comparisons to controls. (**C, D**) Locomotor activity and cumulative locomotor responses (**C**^**1**^ and **D**^**1**^) of animals treated with the DAT blocker GBR during the anticipatory phase (**C, C**^**1**^) or 1-hour after intermittent access to water (Ctr+GBR) or palatable diet (Binge+GBR) (**D, D**^**1**^). Results are expressed as beam breaks (bb). Statistics: **p<0.01 Binge+GBR *vs* Control+GBR. (**E**) Palatable food intake after vehicle (Veh+Binge) or D1R antagonist SCH23390 (SCH+Binge) pretreatment. Note: SCH23390 was acutely administered 30 min before binge session. Statistics: ***p<0.001 SCH+Binge *vs* Veh+Binge. (**F**) Palatable food intake after vehicle (Veh+Binge) or D2R antagonist haloperidol 0.25 mg/kg or 0.5 mg/kg (H^0.25^+Binge and H^0.5^+Binge) treatment. Note: haloperidol was acutely administered 30 min before binge session. (**G**) Immunolabeling of P-S6 in the DS and NAc (for images see also **Suppl. Fig. 2E**) and their associated quantifications (**G**^**1**^, **G**^**2**^, **H**^**1**^, **H**^**2**^) in mice pretreated with SCH23390 or vehicle and exposed to time-locked palatable diet. Scale bar: 50 μm. Statistics: ***p<0.001 Veh+Binge *vs* Veh+Control, ^###^p<0.001 SCH+Binge *vs* Veh+Binge. (**I**) Locomotor activity and cumulative locomotor responses (**I**^**1**^) of animals receiving SCH (SCH+Binge) or vehicle (Veh+Binge) (red arrow) with access to palatable diet (black arrow). Statistics: **p<0.01 SCH+Binge *vs* Veh+Binge. (**J**) Cumulative regular chow intake following acute SCH23390 (SCH+Binge) or vehicle (Veh+Binge). Statistics: **p<0.01 SCH+Binge *vs* Veh+Binge. (**K**) Locomotor activity and cumulative locomotor responses (2 hrs and 30 min, **K**^**1**^) induced by the D1R agonist SKF81297 administered 1 hour after access to time-locked water (Ctr+SKF) or palatable diet (Binge+SKF). Statistics: *p<0.05 and **p<0.01 Binge+SKF *vs* Control+SKF. For number of mice/group and statistical details see **Suppl. Table 1**.

Next, we wondered whether food-reward anticipatory and/or consummatory phases were followed by adaptive changes in DA signaling. We treated mice with GBR12909 (10 mg/kg), a specific DA transporter (DAT) blocker that leads to striatal/accumbal synaptic accumulation of DA. Interestingly, we observed a different behavior depending on BE phases (anticipatory *vs* consummatory). Before palatable food access, GBR similarly increased locomotor activity in both bingeing and control animals (**Fig. 2C, C^1^**). However, when GBR was administered following palatable food consumption (1h), GBR-induced locomotor response was blunted in bingeing animals (**Fig. 2D, D^1^**). These results indicate that BE-induced physiological adaptations are characterized by the enabled ability for palatable food to impinge on DA release and action.

At the postsynaptic level, DA acts onto medium spiny neurons (MSNs) which express either the D1R or D2R. To discriminate their roles in BE, we pretreated animals with the D1R antagonist SCH23390 (0.1 mg/kg) or vehicle (Veh) 30 min prior access to palatable diet. SCH23390 dramatically reduced palatable food consumption (**Fig. 2E**). On the contrary, 30 min pretreatment with the D2R antagonist haloperidol (0.25 and 0.5 mg/kg) did not dampen palatable food consumption (**Fig. 2F**), even at cataleptic doses [30]. In line with this evidence, activation of D1R leads to downstream phosphorylation of S6 and ERK [28, 29]. The adaptive molecular changes also required D1R activation since SCH23990 (0.1 mg/kg) largely suppressed BE-induced P-S6 in both DS (**Fig. 2G, G^1^, H^1^**) and NAc (**Fig. 2G^2^, H^2^**, **Suppl. Fig. 2F**). Of note, although SCH23390 reduced BE-elicited anticipatory locomotor activity, basal locomotor activity in naive animals was not impaired (**Fig. 2I, I^1^**), excluding the confounding effects due to changes in basal locomotor activity. Furthermore, a compensatory rescue in chow intake was observed in SCH23390-pretreated bingeing animals during the dark phase (**Fig. 2J**), excluding potential long-lasting effects of the D1R inhibition. To further validate the implication of D1R in BE-elicited DA modifications, we measured the locomotor activity triggered by the D1R agonist SKF81297 (5 mg/kg) at the end of the BE session. Interestingly, we observed an earlier (first 30 min) significant increase in locomotor activity in bingeing animals compared to control mice, although no major differences were detected during the cumulative 2-hrs response (**Fig. 2K, K^1^**).

Overall, our results reveal that the critical phases surrounding palatable food consumption in the context of BE profoundly affect D1R-associated signaling.

### Peripheral endocannabinoids govern binge eating and metabolic efficiency

Recent studies have highlighted the role of neuronal and endocrine gut systems in the regulation of food reward-seeking and DA-associated behaviors [31, 32]. We therefore tested whether gut-born metabolic signals had a privileged action onto BE when compared to other known circulating satiety signals.

First, we observed that peripherally injected leptin (repeated 0.25 mg/kg, **Suppl. Fig. 3A, B**) or insulin (acute, 0.5 U/kg) did not trigger any reduction in palatable food consumption when administered in bingeing animals (**Fig. 3A**). Then, we investigated whether gut-born satiety signals retained anorectic properties with a similar protocol. GLP-1R agonists, exendin-4 (10 µg/kg) and liraglutide (100 µg/kg), successfully reduced binge-consumption of palatable diet (**Fig. 3A**). Moreover, pharmacological activation of GLP-1R, which did not alter spontaneous ambulatory activity (**Suppl. Fig. 3C**), was also associated with a decrease in the anticipatory and consummatory locomotor phases (**Suppl. Fig. 3D**). Similarly, the cholecystokinin analog CCK-8S (10 µg/kg) acutely decreased palatable food intake (**Fig. 3A**). Since only the anorectic action of gut-born signals was efficient in counteracting binge-like consumption, we also investigated the effect of endocannabinoids (eCBs) which are important mediators of nutrients-induced adaptive responses within the gut-brain axis [33, 34]. We acutely inhibited the CB1R with the global acting antagonist/inverse agonist AM251 (3 mg/kg, i.p.) and observed a dramatic reduction of BE-like consumption (**Fig. 3A**).

**Fig. 3:**
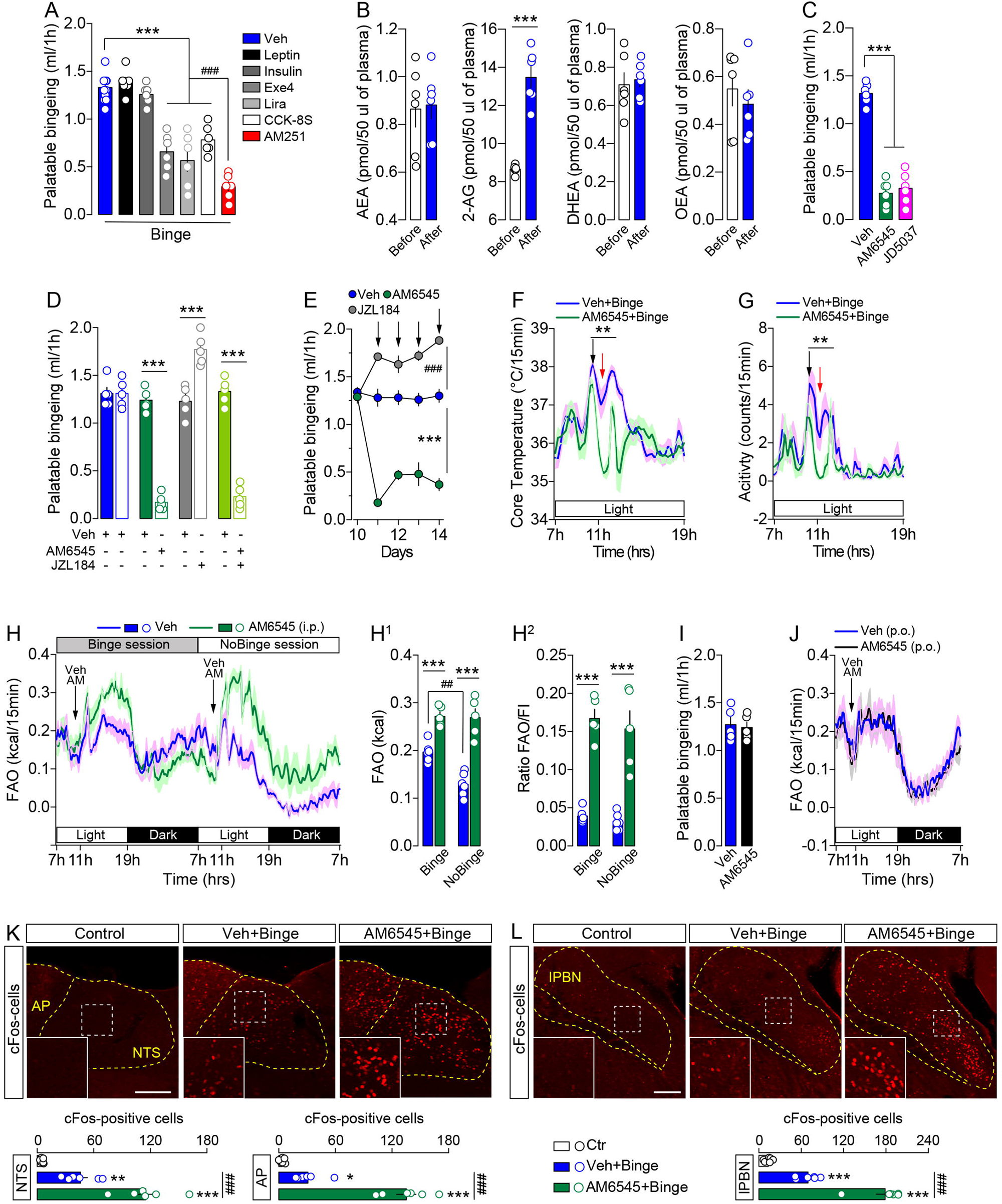
Peripheral endocannabinoids (eCBs) govern binge eating. (**A**) Bingeing in animals pretreated (1h prior binge session) with vehicle (Veh), leptin, insulin, GLP1 agonists exendin-4 (Exe4) and liraglutide (Lira), CCK octapeptide sulfated (CCK-8S) or CB1R inverse agonist AM251. Aside leptin (2 injections/day for 2 consecutive days, **Suppl. Fig. 3A, B**), drugs were administered acutely. Statistics: ***p<0.001 Exe4-, Lira-, CCK-8S-& AM251-treated bingeing mice *vs* Veh+Binge mice, ^###^p<0.001 AM251-treated *vs* Exe4-, Lira & CCK-8S-treated bingeing mice. (**B**) Dosage of peripheral and circulating eCBs: anandamide (AEA), diacylglycerol (2-AG), docosahexanoyl ethanolamide (DHEA) and oleoylethanolamide (OEA) 1 hour before and after palatable bingeing. (**C**) Palatable bingeing in mice (3 males and females/group) pre-treated with a single i.p. injection of vehicle (Veh), peripheral CB1R antagonist AM6545 (10 mg/kg) and peripheral CB1R inverse agonist JD-5037 (3 mg/kg). Statistics: ***p<0.001 AM6545 and JD-5037 *vs* Veh-Binge. (**D**) Palatable bingeing in mice pre-treated with a single i.p. injection of vehicle (Veh), peripheral CB1R antagonist AM6545 (10 mg/kg), and/or MAGL inhibitor JZL184 (8 mg/kg). Statistics: ***p<0.001 AM6545, JZL184, AM6545+JZL184 *vs* Veh-Binge. (**E**) Chronic (4 days) administration of JZL184 and AM6545 on palatable bingeing. Statistics: ***p<0.001 AM6545-Binge *vs* Veh-Binge, ^###^p<0.001 JZL184-Binge *vs* Veh-Binge. (**F**) Effects of acute AM6545 on core temperature. Statistics: **p<0.01 AM6545-Binge *vs* Veh-Binge. Note: black and red arrows indicate administration of AM6545 and palatable food access, respectively. (**G**) Effects of acute AM6545 on locomotor activity matching the core temperature measurements. Statistics: **p<0.001 AM6545+Binge *vs* Veh+Binge. (**H**) Longitudinal measurement of fatty acid oxidation (FAO) following administration of AM6545 during Binge and NoBinge sessions. (**H**^**1**^) Averaged FAO from time of injection (11h00) till the end of light phase (19h00). (**H**^**2**^) Ratio of FAO and food intake (FI) to discriminate between the effect of AM6545 and calories intake. Statistics: ***p<0.001 AM6545 *vs* Veh (in both Binge and NoBinge sessions). (**I**) Bingeing after acute oral gavage of AM6545 (10 mg/kg, p.o.) and (**J**) associated FAO. (**K, L**) Immunolabeling and quantifications of cFos in the cNTS/AP regions (**K**) and in the lPBN (**L**) of control or bingeing animals (males and females) acutely pretreated with Veh or AM6545. Scale bars: 250 μm. Statistics: ***p<0.001, **p<0.01 and *p<0.005 Veh+Binge or AM6545+Binge *vs* Ctr, ^###^p<0.001 AM6545+Binge *vs* Veh+Binge. For number of mice/group and statistical details see **Suppl. Table 1**.

Next, we wondered whether bingeing was accompanied by alterations in circulating peripheral eCBs [anandamide (AEA) and 2-arachidonoylglycerol (2-AG)] and eCBs-related species [docosahexanoyl ethanolamide (DHEA), oleoylethanolamide (OEA)]. While circulating *N*-acylethanolamines (AEA, DHEA, OEA) remained unaffected, bingeing induced a significant increase in circulating 2-AG (**Fig. 3B**). No differences in 2-AG levels were detected in key brain regions such as the hypothalamus, VTA, DS and NAc (**Suppl. Fig. 4A-D**).

Given the rise in peripheral 2-AG, we were eager to explore the role of peripheral CB1R signaling in BE outputs. Thus, we used the *bona fide* peripherally restricted CB1R neutral antagonist AM6545 (10 mg/kg, i.p.) or inverse agonist JD-5037 (3 mg/kg, i.p.), two compounds with poor brain permeability [35–37]. Pretreatment (1h before bingeing) with AM6545 or JD-5037 induced a stark reduction of BE consumption when acutely administered (**Fig. 3C**). Conversely, the increase of 2-AG, achieved through the pharmacological inhibition (JZL184, 8 mg/kg) of its catabolic enzyme monoacylglycerol lipase (MAGL) [38], resulted in an increase of palatable food consumption that was fully prevented by AM6545 (**Fig. 3D**). This bidirectional modulatory action of eCBs/CB1R on BE did not show signs of desensitization and remained efficient throughout 4 days of daily pharmacological intervention (**Fig. 3E**). In the same line, thermogenic and locomotor activity analyses revealed that acute pretreatment with AM6545 strongly dampened both the anticipatory and consummatory phases of BE (**Fig. 3F, G**).

We next explored whether and how peripheral CB1R signaling modulates metabolic efficiency in the context of BE. Pretreatment with AM6545 significantly increased fatty acid oxidation (FAO) (**Fig. 3H, H^1^**) which did not depend on reduced calorie intake (Binge session) or basal calorie contents (NoBinge session) (**Fig. 3H^2^**) nor on altered energy expenditure (EE) (**Suppl. Fig. 4E**). These results indicate that acute manipulation of peripheral eCB tone affects nutrients partitioning and promotes a shift towards whole-body lipid-substrate utilization. Interestingly, oral administration of AM6545 did not blunt bingeing responses (**Fig. 3I**) nor increase FAO (**Fig. 3J**). This evidence suggests that, in our model, CB1R-mediated homeostatic adaptations may not depend on lumen-oriented apical CB1R of endothelial or enteroendocrine intestinal cells [39, 40] but rather on non-lumen-oriented CB1R.

Given these results, we investigated the anatomo-functional structures at the interface between peripheral and central systems. Compared to control and bingeing mice, AM6545-pretreated bingeing mice showed a more pronounced neuronal activation (cFos) in the caudal nucleus tractus solitarius (cNTS) and the area postrema (AP) (**Fig. 3K**) as well as in the cNTS-projecting lateral parabrachial nucleus (lPBN) (**Fig. 3L**), thus pointing to the gut-brain vagal axis as a potential mediator of our effects.

These results indicate that peripheral CB1R signaling is sufficient to control BE by modulating the activity of satietogenic/homeostatic paths.

### The gut-brain vagal axis is required for eCBs-mediated effects

Recent reports have indicated that CB1R is densely expressed in vagal afferent neurons [41]. To discriminate among vagal afferents, we performed a meta-analysis on single-cell transcriptomic results [9] obtained through a path-specific viral strategy (**Fig. 4A**). This analysis revealed that *Cnr1*, but not *Cnr2*, was enriched in all gut-brain vagal segments (**Fig. 4B, Suppl. Fig. 4F, G**) and that, together with well-known afferent markers (*Slc17a6, Scn10a, Htr3a, Cartpt, Grin1, Phox2b*), *Cnr1* may be considered as a constitutive marker of vagal sensory neurons. Thus, we took advantage of subdiaphragmatic vagotomy (VGX) to investigate whether the eCBs-vagus axis was necessary/sufficient to mediate the modulatory effects of eCBs on BE.

**Fig. 4:**
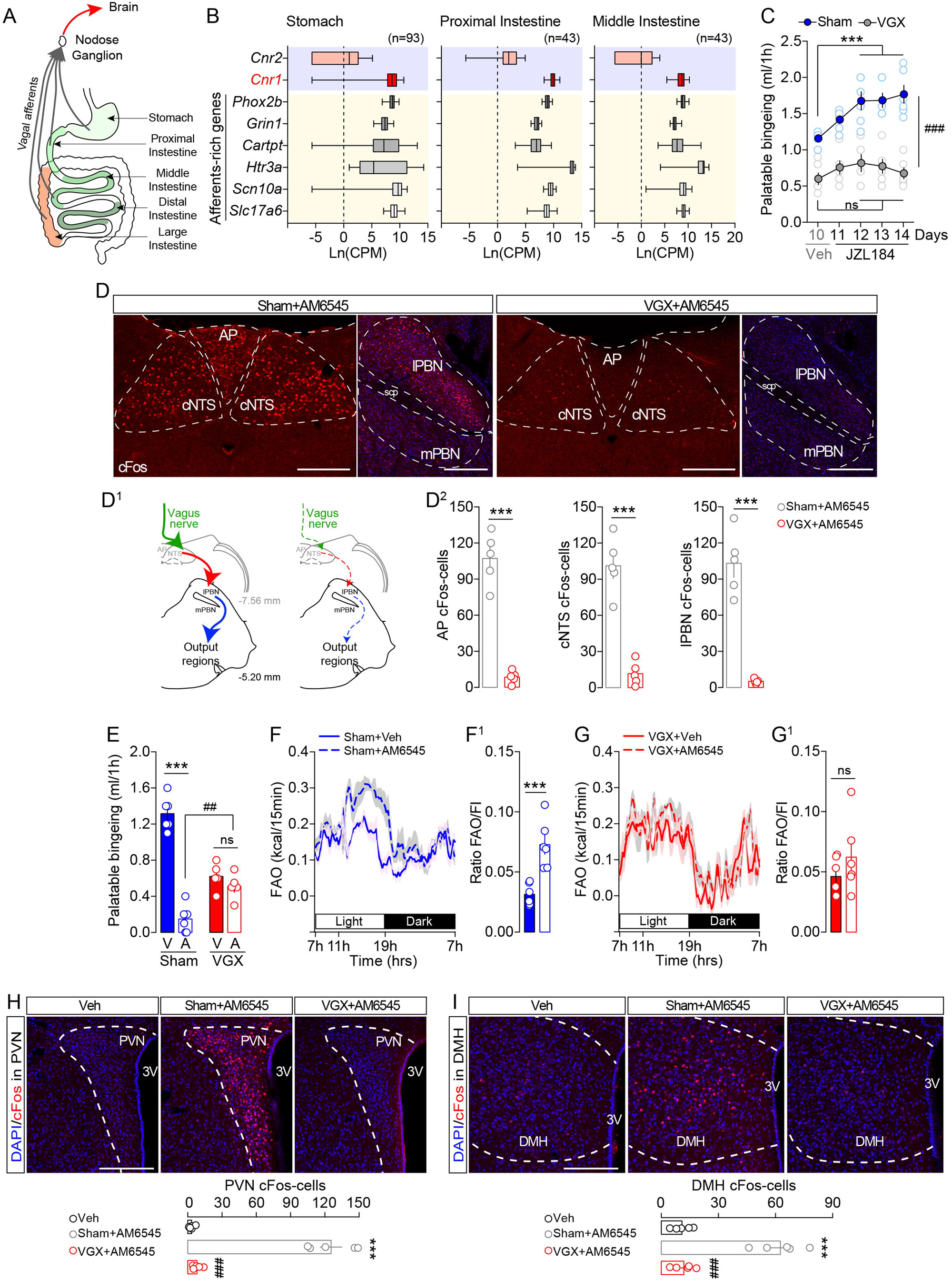
The gut-brain vagal axis is required for eCBs-mediated effects. (**A**) Scheme indicates gut-originated vagal afferents virally targeted for single-cell transcriptomic analysis [9]. (**B**) Enrichment of vagal markers (*Slc17a6, Scn10a, Htr3a, Cartpt, Grin1, Phox2b*) and comparison with *Cnr1* and *Cnr2* in sensory vagal neurons labeled from microinjections in the stomach, proximal and middle intestines. Note: for other gut-brain vagal segments see **Suppl. Fig. 4F, G**. (**C**) Palatable food consumption in sham and vagotomized (VGX) animals pre-treated with Veh (Day 10) or JZL184 (Day 11-14) 2-hrs before bingeing sessions. Statistics: ***p<0.001 Sham+JZL184 *vs* Sham+Veh, ^###^p<0.001 VGX+JZL184 *vs* Sham+JZL184. (**D**) cFos immunolabeling in the area postrema (AP), caudal nucleus tractus solitarius (cNTS), lateral parabrachial nucleus (lPBN) and medial parabrachial nucleus (mPBN) in sham and vagotomized animals treated with the peripheral CB1R antagonist AM6545 (10 mg/kg). Scale bars: 250 μm. (**D**^**1**^) Scheme indicates the central vagus→cNTS→PBN→target regions path in sham and VGX mice. (**D**^**2**^) Quantification of cFos-positive neurons in the AP, cNTS and lPBN in sham and VGX mice injected with AM6545. Statistics: ***p<0.001 VGX+AM6545 *vs* Sham+AM6545. (**E**) Palatable bingeing in sham and vagotomized (VGX) animals pre-treated with AM6545 (A) or vehicle (V), and associated measurements of FAO (**F, F**^**1**^ and **G, G**^**1**^). Statistics: ***p<0.001 Sham+AM6545 *vs* Sham+Veh. (**H, I**) cFos immunolabeling in the paraventricular nucleus (PVN) and dorsomedial nucleus of the hypothalamus (DMH) of sham or VGX animals acutely treated with vehicle or AM6545 and associated counting. Scale bars: 250 μm. Statistics: ***p<0.001 Sham+AM6545 *vs* Veh, ^###^p<0.001 VGX+AM6545 *vs* Sham+AM6545. For number of mice/group and statistical details see **Suppl. Table 1**.

First, we tested whether the pro-bingeing effects of the MAGL inhibitor JZL184 (**Fig. 3D, E**) required an intact vagal transmission. Although vagotomy *per se* was associated with a decrease in time-locked palatable feeding (see **Fig. 4C, E**) and consequent BE-derived compensatory homeostatic adaptations (**Suppl. Fig. 5**), repeated inhibition (4 days) of MAGL (JZL184, 8 mg/kg) induced a significant increase of palatable food consumption only in sham animals (**Fig. 4C**).

Second, we tested whether the anti-bingeing effects of AM6545 were also routed by the vagus nerve. In sham mice, acute AM6545 led to a strong increase of cFos in the cNTS, AP and cNTS-projecting lPBN, while the signal was abolished in VGX mice (**Fig. 4D, D^1^, D^2^**). Of note, blockade of peripheral CB1R did not activate the rostral NTS (rNTS) (**Suppl. Fig. 4H**), thus indicating and further confirming that gut-to-brain vagal inputs are necessary to mediate the action of AM6545.

We also observed that vagal integrity was essential to mediate the anorectic action of AM6545 on BE since it did not trigger anti-bingeing responses in VGX mice compared to sham mice (**Fig. 4E**). Furthermore, vagotomy abolished the increase in FAO following AM6545 administration as observed in sham mice (**Fig. 4F, F^1^, G, G^1^**), indicating that the gut-brain vagal communication routes feeding and the metabolic components associated with BE.

These vagus-dependent homeostatic adaptations promoted by the peripheral blockade of CB1R prompted us to investigate whether AM6545 was able to alter the activity of brainstem-projecting hypothalamic structures that control feeding. AM6545 induced a strong vagus-dependent increase of cFos in the PVN and DMH regions (**Fig. 4H, I**), indicating that the metabolic adaptations induced by peripheral blockade of CB1R require a vagus-mediated cNTS→PBN→hypothalamus circuit whose nodes’ activation control feeding and energy homeostasis [42–44].

### Peripheral CB1R signaling routed by the vagus nerve controls the activity of VTA dopamine neurons

Palatable bingeing also strongly relies on central DA-dependent mechanisms (**Fig. 2**). Therefore, we investigated whether blockade of peripheral CB1R was also efficient in counteracting BE-induced molecular adaptations in dopaminoceptive structures (**Fig. 2**). Indeed, AM6545 reduced BE-induced activation of S6 (marker of translational activity) and cFos (marker of neuronal activity) in the DS and NAc (**Fig. 5A**) with no sex-dependent differences (**Suppl. Fig. 6A**).

**Fig. 5:**
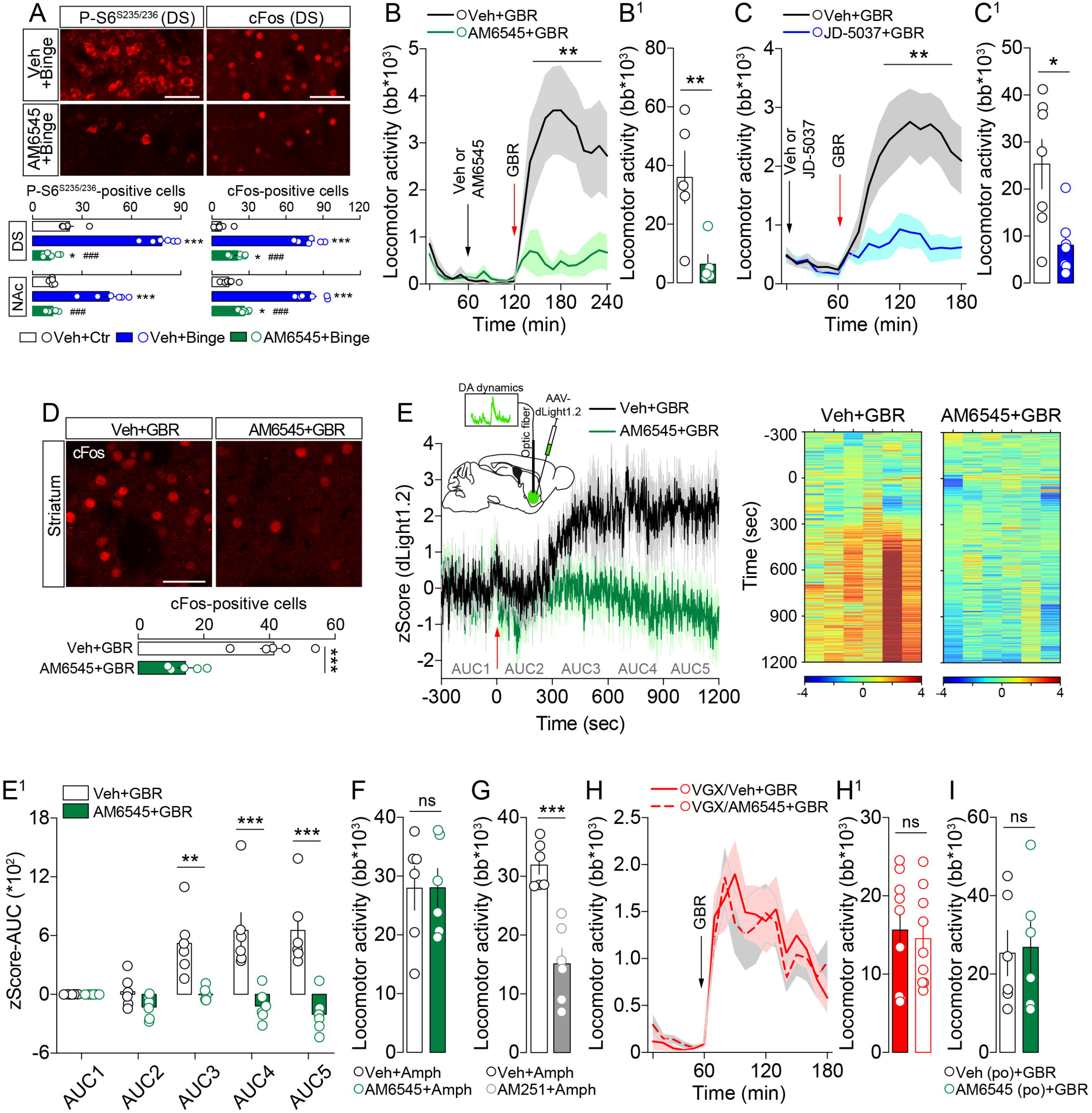
Peripheral CB1R signaling modulates dopamine dynamics. (**A**) Immunolabeling and quantifications of P-S6 and cFos in the DS and NAc of control or bingeing animals (males and females) acutely pretreated with Veh or AM6545. Scale bars: 50 μm. Statistics: ***p<0.001 and *p<0.005 Veh+Binge or AM6545+Binge *vs* Ctr, ^###^p<0.001 AM6545+Binge *vs* Veh+Binge. (**B, B**^**1**^) Effect of AM6545 or Veh on GBR-induced locomotor activity (beam breaks, bb). Statistics: **p<0.01 AM6545+GBR *vs* Veh+GBR. (**C, C**^**1**^) Effect of JD-5037 or Veh on GBR-induced locomotor activity. Statistics: **p<0.01, *p<0.05 AM6545+GBR *vs* Veh+GBR. (**D**) Effect of AM6545 on GBR-triggered striatal cFos expression. Scale bar: 50 μm. Statistics: ***p<0.001 AM6545+GBR *vs* Veh+GBR. (**E**) Longitudinal profile and heat maps of GBR-induced accumbal DA accumulation (dLight1.2) in mice pre-treated with Veh or AM6545 1h before GBR12909 (red arrow). (**E**^**1**^) Quantification of bulk fluorescence (AUC) in Veh+GBR and AM6545+GBR groups. Statistics: ***p<0.001 AM6545+GBR *vs* Veh+GBR. (**F**) Cumulative locomotor activity response in mice pretreated with vehicle (Veh+Amph) or AM6545 (AM6545+Amph). For longitudinal locomotor activity see **Suppl. Fig. 6B**. (**G**) Cumulative locomotor activity response in mice pretreated with vehicle (Veh+Amph) or AM251 (AM251+Amph). Statistics: ***p<0.001 AM251+Amph *vs* Veh+Amph. (**H**) GBR-induced locomotor activity and cumulative locomotor response (**H**^**1**^) in VGX mice pretreated with vehicle (VGX/Veh+GBR) or AM6545 (VGX/AM6545+GBR). (**I**) Cumulative locomotor response in mice pretreated with oral gavage (po) of vehicle (Veh (po)+GBR) or AM6545 (AM6545 (po)+GBR). For longitudinal locomotor activity (**Suppl. Fig. 6C**). For number of mice/group and statistical details see **Suppl. Table 1**.

Next, we explored the functional connection between peripheral eCBs and the gut-to-brain vagal axis in the modulation of the DA system. Since hedonic/motivated spontaneous feeding is blunted in AM6545-treated mice (**Fig. 3, 4**), thus limiting further *in vivo* investigations of DA/reward events, we promoted and gauged the activity of the DA system by using pharmacological and behavioral strategies.

Naive mice were acutely pretreated with AM6545 (or JD-5037) or vehicle 1h before being administered with the DAT blocker GBR12909. Blockade of peripheral CB1R drastically reduced GBR-induced locomotor activity (**Fig. 5B, B^1^, C, C^1^**) as well as GBR-triggered striatal cFos (**Fig. 5D, D^1^**). To further study *in vivo* DA dynamics, we took advantage of the virally expressed DA biosensor dLight1.2 coupled to *in vivo* fiber photometry [22]. Importantly, pretreatment with AM6545 blunted GBR-evoked accumbal DA accumulation (**Fig. 5E, E^1^**). Interestingly, unlike the brain penetrant CB1R antagonist/inverse agonist AM251, the peripheral AM6545 failed in counteracting amphetamine-induced locomotor activity (**Fig. 5F, G, Suppl. Fig. 6B**). This evidence indicates a clear difference in the action of peripheral *vs* central CB1R and suggests that inhibition of peripheral CB1R may modulate the intrinsic spontaneous activity of DA-neurons rather than altering evoked DA release events. Altogether, these results reveal that inhibition of peripheral CB1R, besides promoting satiety and FAO (**Fig. 3, 4**), may dampen reward-driven feeding also by concomitantly reducing DA-neurons spontaneous activity and consequent activation of dopaminoceptive structures.

To directly address this point, VGX mice were acutely pretreated with AM6545 1h prior GBR12909. Remarkably, ablation of the vagus nerve abolished the blunting effect of AM6545 on GBR-elicited locomotor activity (**Fig. 5H, H^1^**). Moreover, consistently with what observed for palatable bingeing (**Fig. 3I**), this vagus-to-brain effect was further emphasized by the lack of action of AM6545 when orally administered (**Fig. 5I, Suppl. Fig. 6C**).

Finally, to fully establish that peripheral inhibition of CB1R modulates the somatic activity of VTA DA-neurons, we performed cell type-specific *in vivo* Ca^2+^ imaging of DA-neurons in presence or absence of AM6545. We took advantage of *Drd2*-Cre mice to virally express GCaMP6f in VTA DA-neurons (**Fig. 6A**) as they co-express the autoreceptor D2R (**Suppl. Fig. 6D**). Using this strategy, we were able to detect activation and inhibition of VTA DA-neurons following rewarding or aversive events (**Suppl. Fig. 6E, F**). To avoid the above-mentioned *in vivo* limiting factors (hypophagia under AM6545) and the confounding effects of the D2R-dependent autoinhibition of psychostimulants-evoked local (VTA) somatodendritic DA accumulation, we used two behavioral paradigms that readily and transiently modulate DA-neurons activity: the tail suspension and exposure to a new environment (**Fig. 6B-D**). Blockade of peripheral CB1R led to a reduced activation of VTA DA-neurons in both paradigms (**Fig. 6C, D**). Behaviorally, while AM6545 did not alter the time of immobility in the tail suspension test (**Fig. 6E**), it reduced novelty-induced exploratory drive in a vagus-dependent manner (**Fig. 6F, F^1^**). Of note, the brain permeable AM251 reduced exploratory drive in a vagus-independent manner (**Suppl. Fig. 6G**).

**Fig. 6:**
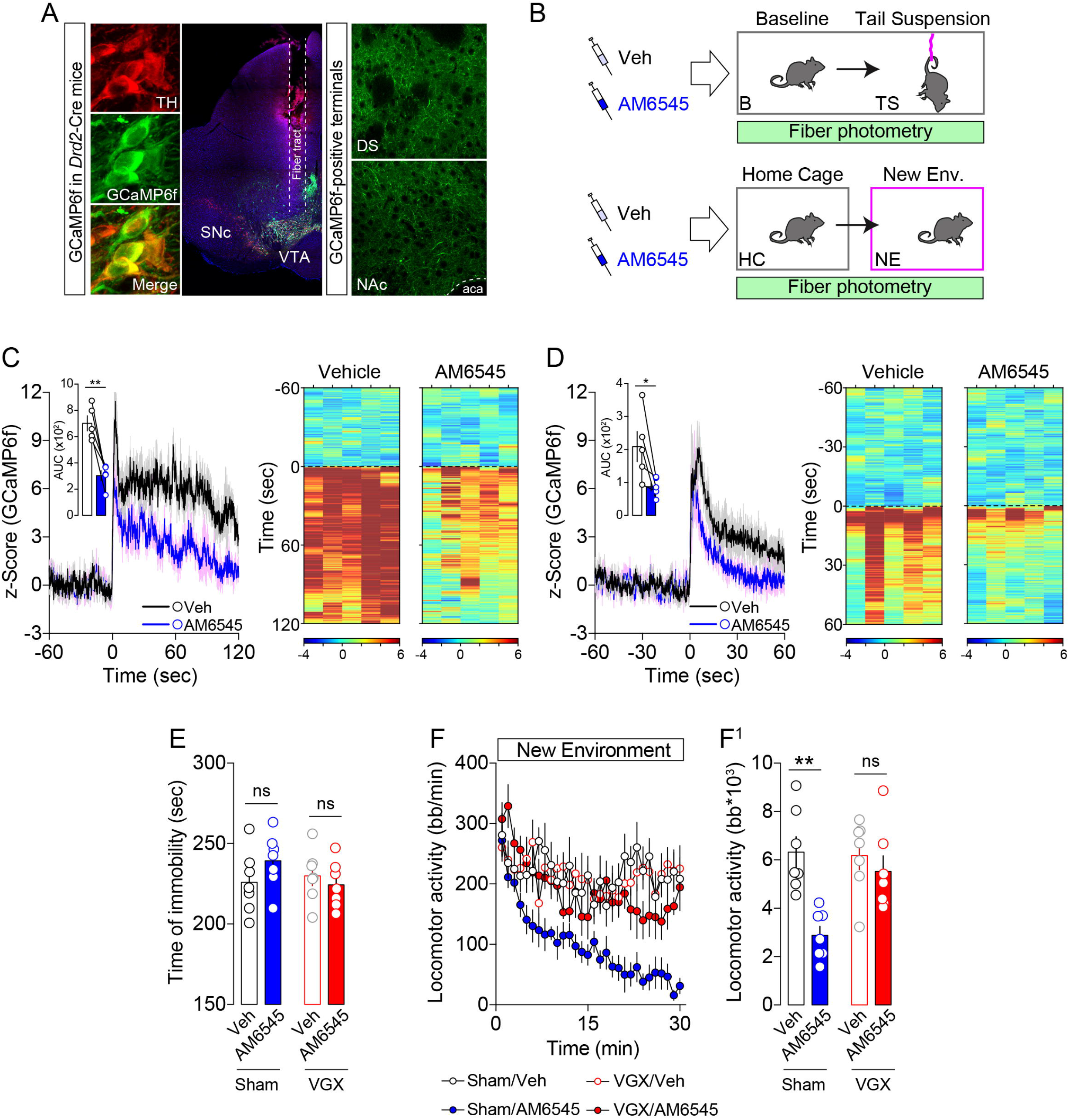
Peripheral CB1R signaling routed by the vagus nerve controls VTA DA-neurons activity. (**A**) Expression of GCaMP6f in VTA DA-neurons of virally injected *Drd2*-Cre mice. Note (*i*) colocalization with TH and GCaMP6f-positive neurons and (*ii*) projecting terminals in the DS and NAc. See also, **Suppl. Fig. 6D** for TH expression in the VTA of *Drd2*-eGFP mice. (**B**) Behavioral paradigms used to trigger the activity of VTA DA-neurons: tail suspension and exposure to a new environment (NE). [For validation of *in vivo* recording of Ca^2+^ transients in VTA D2R-(DA)-neurons see **Suppl. Fig. 6E, F]**. (**C, D**) Temporal dynamics and corresponding heat maps of DA-neurons activity during the tail suspension test (**C**) and exposure to a new environment (**D**). Statistics: *p<0.05, **p<0.01 AM6545 *vs* Veh. (**E**) Immobility time (sec) of sham and VGX mice acutely pretreated with Veh or AM6545 1 hour before tail suspension (6 min). (**F**) Effect of AM6545 or Veh in sham and VGX mice on novel environment-induced locomotor activity. (**F**^**1**^) Cumulative locomotor responses in sham and VGX mice pretreated with vehicle or AM6545. Statistics: **p<0.01 Sham/AM6545 *vs* Sham/Veh. For number of mice/group and statistical details see **Suppl. Table 1**.

Collectively, these results reveal that peripheral CB1R signaling routed through the vagal axis exerts an integrative control over metabolic/satietogenic (**Fig. 3, 4**) and DA paths (**Fig. 5, 6**), both of which contribute to the establishment of BE.

## Discussion

A characteristic feature of feeding behavior is its key ability to dynamically adapt to sensory and environmental stimuli signaling food availability. Such adaptive strategy is even more pronounced when food is palatable and energy-dense. Indeed, the control of feeding strategies requires complex and highly interactive systems that can hardly be unequivocally attributed to single structures, circuits or mediators.

In our study, we observed that, first, palatable time-locked feeding mobilizes both homeostatic and hedonic components of feeding through fast, but yet physiological, allostatic adaptations. Second, such allostatic adaptations require a concerted involvement of central DA (hedonic drive) and peripheral eCBs signaling (homeostatic and hedonic drive). Third, the permissive role of peripheral eCBs fully relies on the vagus nerve which, by a polysynaptic circuit, controls the activity of both satietogenic and reward (DA) structures. Fourth, our results point to peripheral CB1R as promising therapeutic targets to counteract eating as well as reward-related disorders.

Overall, our study describes for the first time the fundamental role of eCB gut-brain transmission as a core component of binge eating and its behavioral, cellular and molecular adaptations.

Here, by investigating the pathways involved in hedonic feeding in absence of hunger or energy deprivation, we provide evidence that the hedonic drive to eat, as triggered by our intermittent time-locked model, promotes rapid homeostatic compensations leading to escalating consumption of palatable food and to allostatic adaptations of energy metabolism. As such, caloric demands are fulfilled and classical energy-mediated homeostatic signals (leptin, insulin) do not seem to spontaneously interfere, thus providing us the opportunity to study food intake-related integrative pathways with the abstraction of the homeostatic *vs* hedonic discrepancy. In line with clinical data [45, 46], we observed that binge-like feeding in lean animals is not necessarily associated with overweight gain. The allostatic adaptations observed, ranging from increased anticipatory feeding phase to pre-feeding increased corticosterone levels and food intake maximization, all represent key hallmarks of the compulsive and emotional states of BE patients [47–49]. The anticipatory feeding phase was associated with decreased levels of plasma TG and insulin, whereas both anticipatory and consummatory phases were characterized by increased energy expenditure, core temperature and metabolic efficiency, thereby suggesting a metabolic shift of nutrients’ use. This observation perfectly mirrors the allostatic theory, which stands on the fact that an organism anticipates and adapts to environmental changes while accordingly adjusting several physiological parameters to maintain stable homeostatic states [50, 51]. Allostatic mechanisms have been classically discussed in terms of stress-related regulatory events. However, the hedonic value of a stimulus (food, recreational drugs) can function as a feed-forward allostatic factor [4].

In line with this notion, analysis of key DA-activated downstream targets in the DS and NAc highlighted specific patterns of molecular activation. Notably, while the anticipatory phase was associated with an increase in P-ERK, the consummatory phase was also accompanied by a robust increase in P-S6. Such signaling events, which did not depend on a single episode of palatable food intake, required the dopamine D1R. Whether this molecular regulation reflects the full establishment of BE or the amount of ingested palatable food remains to be established. Nevertheless, this D1R mechanism is of interest since, contrary to the well-known molecular insights of drugs of abuse also requiring the D1R [52–54], food-related disorders have been predominantly associated with altered D2R signaling [55, 56]. Our results reveal that binge eating, characterized by transients and sudden urges of hedonic drive, requires, at least in its early phases, a D1R-mediated transmission. This D1R-dependent mechanism is in line with the affinity and time-dependent dynamics of dopamine effects [52] as well as with the molecular action of released DA which, by binding to Gα(olf)-coupled D1R, would trigger the activation of the aforementioned pathways, while activation of the Gi-coupled D2R would lead to their inhibition. However, in clear opposition to psychostimulants, which directly act at central DA synapses, food and food-mediated behaviors impact on DA transmission through a plethora of indirect and often peripherally born long-range acting mediators. Notably, nutrients, as demonstrated by intragastric infusion of fat and sugar [14, 57–59], or gut-born signals [60–62], are sufficient to modulate DA release in reward-related structures. Here, we observed that gut-born signals such as CCK, GLP1 and endocannabinoids (eCBs) are essential in gating bingeing. In particular, we found that time-locked consumption of palatable food was associated with a rise in peripheral eCBs, notably 2-AG. Furthermore, inhibition of the 2-AG-degrading MAGL resulted in a potentiation of palatable food consumption. Thus, by taking advantage of low brain permeant CB1R antagonists/inverse agonists, we observed that blockade of CB1R was able to fully abolish both anticipatory and consummatory phases of hedonic feeding as well as the potentiated feeding induced by MAGL inhibition. These effects agree with the literature showing that peripheral eCBs are highly and dynamically modulated in eating disorders, and act as powerful mediators of the gut-to-brain integration [17].

Previous studies have shown that chronic administration of AM6545 promoted long-term maintenance of weight loss and reduction of dyslipidemia in obesity [35, 37]. Here, we show that a single, as well as repeated, administration of AM6545 potently inhibits binge eating and its molecular adaptations. The anorectic effects of peripheral blockade of CB1R have been attributed, at least in part, to the property of global CB1R antagonists to promote fatty acid oxidation (FAO). In agreement with these studies, we have observed that acute administration of AM6545 was able to dramatically increase FAO. However, here we also demonstrate that such effects require the vagus nerve. The action of eCBs as well as of AM6545 on CB1R-expressing vagal afferents [41] may explain our results. In fact, an increase in eCBs during palatable feeding would slow the vagus nerve activity through the inhibitory Gi-coupled signaling of CB1R, thus delaying cNTS-reaching satiety signals and promoting food intake. On the contrary, peripheral blockade of CB1R, especially when peripheral eCB levels are endogenously high (*i*.*e*. binge eating, bulimia, obesity), would lead to a prompt disinhibition and to the concomitant activation of satietogenic brain pathways (cNTS→PBN→PVN). Moreover, it is worth to mention that under fasting or lipoprivic conditions the systemic CB1R inverse agonist SR141716A modulated feeding by the vagal and sympathetic systems [63]. Another site of action for peripheral eCBs is represented by CB1R-expressing gut cells [40, 64]. Interestingly, oral administration of a peripheral CB1R antagonist resulted in a reduction of alcohol intake via a ghrelin-dependent and vagus-mediated mechanism [64]. However, in our reward-driven feeding model, oral administration of AM6545 failed to modulate metabolic efficiency as well as to prevent bingeing behavior, thus suggesting that lumen-oriented apical CB1R may not be involved in our mechanism. Intriguingly, recent studies have uncovered that sensory neuropod cells in the gut [65] can synaptically signal with the juxtaposed vagal afferents using, among other possible mediators [66], the fast-acting neurotransmitter glutamate [10]. Whether this specialized gut-to-vagus synapse also mobilizes eCBs, as it occurs at most central excitatory synapses, remains to be determined. In addition, it is important to highlight the key role of peripheral CB1R in adipocytes in the regulation of energy balance [67]. However, whether and how adipocytes may, indeed indirectly, influence the vagal axis is yet unclear.

Overall, it would not be hazardous to suggest that peripheral eCBs may impact feeding patterns through different integrative mechanisms which, depending on the location of peripheral CB1R, may strongly modulate distinct hedonic and homeostatic functional outputs. These results call for cell type- and tissue type-specific strategies to selectively delete CB1R and/or eCBs-producing enzymes in distinct compartments of the gastrointestinal tract and in the neuronal gut-brain axis.

To anatomically provide an explanatory gut-to-brain circuit able to support the vagus-mediated action of AM6545, we found a stark increase of cFos in key brain regions of the satietogenic pathway. Importantly, we reveal that blockade of peripheral CB1R signaling resulted in a strong vagus-dependent activation of the cNTS as well as of its downstream projecting structures, notably the lPBN and the hypothalamic PVN. This segmented activation of the gut→brainstem→hypothalamus path is most likely responsible for the AM6545-induced effects on bingeing and energy homeostasis since activation of these nodes has been shown to reduce food intake and alter energy homeostasis [43, 68–70]. In addition to this satietogenic path and given the strong reward component of our paradigm, we also reveal that AM6545-mediated vagus activation results in a dampened activation of VTA DA-neurons. However, such effect did not depend on the releasing capabilities of DA-neurons since AM6545 failed to alter amphetamine-evoked locomotor activity. In addition, taking advantage of virally mediated GCaMP6f-mediated *in vivo* Ca^2+^ imaging of putative VTA DA-neurons, here we demonstrate that peripheral blockade of CB1R clearly reduces the evoked activity of DA-neurons, a feature resembling some neurochemical effects of vagal nerve stimulation [71, 72].

The VTA is characterized by a highly heterogeneous connectivity [73], and a single and monosynaptic circuit responsible for the inhibition of DA-neurons through the AM6545-activated vagus nerve cannot be selectively sorted out yet. However, several satiety-related structures in the brainstem and hypothalamus are known to project and modulate, directly and/or indirectly, VTA DA-neurons [13, 74–77]. Among these circuits, the PBN→VTA relay is of particular interest since excitatory PBN neurons also largely contact VTA GABA-neurons [76, 78] which in turn may drive the inhibition of VTA DA-neurons and consequent dampening of motivated behaviors.

Here, we show that DA-dependent adaptations require orchestrated inputs among which peripheral eCBs, through the vagus nerve, allostatically scale the homeostatic and hedonic components of feeding and act as mandatory gatekeepers for adaptive responses of the reward circuit. The gut-brain axis is increasingly incriminated as a key player of the regulation of energy metabolism [79], and we show that BE is under the control of the vagus-mediated peripheral inputs. Pointing to peripheral eCBs as permissive actors of this eating disorder certainly brings novelty to the clinical investigations aimed at identifying innovative and non-invasive therapeutic strategies. Importantly, this study further points to the gut-brain axis as a privileged target to modulate brain structures that are functionally responsible for processing cognitive and reward events in an integrative manner. In conclusion, while further studies are warranted to fully untangle the key enteric actors responsible for this phenomenon, our study identifies a novel integrative mechanism by which peripheral endocannabinoids through the gut-brain vagal axis gate allostatic feeding by controlling satiety and reward events, thus also paving the way to target peripheral elements to counteract brain disorders.

## Acknowledgments

We thank Chloé Morel, Rim Hassouna, Anne-Sophie Delbes, Daniela Herrera Moro and Raphaël Denis for technical advice and support. Adrien Paquot (BPBL/UCLouvain) is acknowledged for his help with eCB quantification. We thank Olja Kacanski for administrative support, Isabelle Le Parco, Ludovic Maingault, Angélique Dauvin, Aurélie Djemat, Florianne Michel, Magguy Boa and Daniel Quintas for animals’ care. We acknowledge the technical platform Functional and Physiological Exploration platform (FPE) of the Université de Paris (BFA-UMR 8251) and the animal facility Buffon of the Institut Jacques Monod. This work was supported by the Fyssen Foundation, Nutricia Research Foundation, Allen Foundation Inc., *Agence Nationale de la Recherche* (ANR-21-CE14-0021-01), Université de Paris and CNRS. CB and EM were supported by fellowships from the *Fondation pour la Recherche Médicale* (FRM). Telemetry experiments were supported by the Continuous Glucose Telemetry Award 2018 (Dr. Denis) and sponsored by Data Sciences International.

## Author Contributions

C.B. and G.G. conceived, performed and analyzed most of the experiments. J.C. performed surgeries and behavioral experiments. E.M. helped with molecular studies. E.F. performed vagotomy. C.M. helped with fiber photometry. G.G.M. and R.T. analyzed eCB levels. S.L. provided critical feedback. G.G. supervised the whole project and wrote the manuscript with contribution from all coauthors.

