## Supplementary material for "Identification of an endocannabinoid gut-brain vagal mechanism controlling food reward and energy homeostasis": Suppl. Material

Running title: Peripheral endocannabinoids gate reward feeding and homeostasis

Key words: binge eating, dopamine, 2-AG, vagus nerve, striatum, reward, metabolism

**Supplementary material and methods**

**Animals**

Animals were randomly assigned to each experimental group or treatment. When undergoing pharmacological manipulations mice were used once. Only *Drd2*-Cre::GCaMP6f^VTA^ mice as well as sham and VGX C57BL6 mice of **Fig. 6** were used twice in both tail suspension and novelty-induced exploratory drive paradigms. In these cases, mice were allowed to rest for 7-10 days between experiments.

After each experimental design, animals were either sacrificed to investigate molecular (cFos, phospho-ERK, phospho-S6), chemical (endocannabinoid species) and hormonal adaptations, or euthanized.

To minimize stress, animals were habituated to handling and saline injections during 3-4 consecutive days before each experiment.

**Behaviors**

*Palatable binge eating-like paradigm.* Binge-eating protocol lasted till mice reached a stable consumption/maximization of palatable mixture during at least 3 consecutive days. Behavioral and pharmacological experiments in bingeing mice were performed after stable food-reward consumption.

*Tail suspension*. The tail suspension test was performed as previously described [1]. Behavioral analysis (sham and VGX C57BL6 mice pretreated with Veh or AM6545) was performed during a 6-min test. Videos were recorded and blindly scored by two persons. For *in vivo* Ca^2+^ imaging in the VTA of *Drd2*-Cre mice, fiber photometry was performed during the struggling phase (first 2 min).

*HFHS-induced increased VTA activity*. Animals were provided with a high-fat high-sugar (HFHS) pellet to validate the recording of VTA DA-neurons (activation) in *Drd2*-Cre mice. Ca^2+^ imaging acquisition and analysis were performed before and during feeding.

*Scruff restraint inhibits VTA activity*. Animals were immobilized by restraining to validate the recording of VTA DA-neurons (inhibition) in *Drd2*-Cre mice. Ca^2+^ imaging acquisition and analysis were performed before and during scruff restraint.

**Drug treatments**

All drugs and vehicle solution were freshly prepared in the morning immediately before their administration and administered at a body volume of 10 ml/kg.

*Doses*. For pharmacological interventions, doses were chosen according to previous reports: insulin (0.5 U/kg, Novo Nordisk, Lot GT67422, [2]), CCK-8S (10 μg/kg, Tocris, #1166, [3]), liraglutide (100 μg/kg, gift from Novo Nordisk, [4]), exendin-4 (10 μg/kg, Tocris, #1933, [3]), leptin (twice/day for 2 days at 0.25 mg/kg, Tocris, #2985, see **Suppl. Fig. 3A, B**), AM251 (3 mg/kg, Tocris, #1117, [5]), AM6545 (10 mg/kg, Tocris, #5443, [6]), JD-5037 (3 mg/kg, MedChemExpress, #HY-18697, [7, 8]) SKF81297 (5 mg/kg, Tocris, #1447, [9]), haloperidol (0.25 and 0.5 mg/kg, Tocris, #0931, [10]), SCH23390 (0.1 mg/kg, Tocris, #0925, [9]), GBR12909 (10 mg/kg, Tocris, #0421, [9]), d-amphetamine sulphate (2 mg/kg, Tocris, #2813, [11]), JZL184 (8 mg/kg, Tocris, #3836, [12]).

*Preparation of drugs*. Aside AM251, AM6545, JD-5037 and JZL184, all compounds were dissolved in saline. AM251, AM6545, JD-5037 and JZL184 were dissolved in a solution containing DMSO, Kolliphor and saline at the following ratio 2:1:97. Matched vehicles (saline or DMSO/Kolliphor/saline) were administered in control animals.

*Timing of administration*. Insulin, leptin, CCK-8S, liraglutide, exendin-4, AM251, AM6545, JD-5037 were administered 1-hour prior bingeing sessions or other paradigms when indicated (GBR-induced locomotor activity, tail suspension test and novelty-induced exploratory drive). JZL184 was administered 2-hours prior bingeing sessions. SCH23390 and haloperidol were administered 30 min prior bingeing sessions.

**Triglycerides, insulin and corticosterone measurements**

Plasma circulating triglycerides (TG) were measured with a quantitative enzymatic measurement (Serum Triglyceride Determination Kit, Sigma-Aldrich, Saint-Louis, USA). Insulin dosage was performed with ELISA kit (mouse ultrasensitive insulin ELISA, ALPCO, Salem, NH, USA). Corticosterone was measured with RIA kit (MP Biomedicals, Orangeburg, NY, USA). All kits were used according to the manufacturer guidelines.

**Oral glucose tolerance test (OGTT)**

Oral glucose tolerance test was performed following the establishment of bingeing. Animals were fasted 6 hours before oral gavage of glucose (2 g/kg). Blood glucose was measured from the vein blood tail using a glucometer (Menarini Diagnotics, Rungis, France) at 0, 15, 30, 45, 60, 90, and 120 min. Blood samples were taken at 0, 15, 30 and 60 min to measure insulin levels.

**Tissue preparation and immunofluorescence**

Slices were incubated overnight or 48-hrs at 4°C with the following primary antibodies: rabbit anti-phospho-S6 Ser^235/236^ (1:1000, Cell Signaling Technology, #2211), rabbit anti-phospho-S6 Ser^240/244^ (1:1000, Cell Signaling Technology, #2215), rabbit anti-cFos (1:1000, Synaptic Systems, #226 003), chicken anti-GFP (1:1000, Abcam, #Ab13970) or mouse anti-TH (1:1000, Millipore, #MAB318). Sections were incubated for 60 min with the following secondary antibodies: anti-chicken Alexa488 (1:1000, Invitrogen), anti-rabbit or anti-mouse Cy3 AffiniPure (1:1000, Jackson Immunoresearch). Images used for quantification were all single confocal sections. The objectives (10X or 20X) and the pinhole setting (1 airy unit) remained unchanged during the acquisition of a series for all images. Photomicrographs were obtained with the following band-pass and long-pass filter settings: GFP (band pass filter: 505–530) and Cy3 (band pass filter: 560–615).

For each brain structure, quantifications were performed on two sections (both hemispheres), and averaged. Structures were selected according the following coordinates (from bregma, in mm): DS/NAc (1.18 to 0.98), PVN (-0.70 to -0.94), DMH (-1.46 to -1.82), PBN (-5.02 to -5.34), rNTS (-6.84 to -7.08), and AP/cNTS (-7.32 to -7.76).

**Western blotting and quantitative RT-PCR**

*Western blotting*. At the end of the binge session, the mouse head was cut and immediately immersed in liquid nitrogen for 3 seconds. The brain was then removed and dissected on ice-cold surface, sonicated in 200 µl (dorsal striatum) and 100 µl (nucleus accumbens) of 1% SDS supplemented with 0.2% phosphatase inhibitors and 1% protease inhibitors, and boiled for 10 minutes. Aliquots (2.5 µl) of the homogenates were used for protein quantification using a BCA kit (BC Assay Protein Quantitation Kit, Interchim Uptima, Montluçon, France). Equal amounts of proteins (10 µg) supplemented with a Laemmli buffer were loaded onto 10% polyacrylamide gels. Proteins were separated by SDS-PAGE and transferred to PVDF membranes (Millipore). The membranes were immunoblotted with the following antibodies: rabbit anti-phospho-Ser^235/236^-S6 (1:1000, Cell Signaling Technology, #2211), rabbit anti-phospho-Ser^240/244^-S6 (1:1000, Cell Signaling Technology, #2215), rabbit anti-phospho-ERK (1:2000, Cell Signaling Technology, #4370), mouse anti-beta-actin (1:5000, Sigma Aldrich, #A1978). Detection was based on HRP-coupled secondary antibody binding using ECL. The secondary antibodies were anti-mouse (1:5000, Dako, #P0260) and anti-rabbit (1:10000, Cell Signaling Technology, #7074). Membranes were imaged using the Amersham Images 680. Quantifications were performed using the ImageJ software.

*Quantitative RT-PCR*. The hypothalamus was quickly dissected out and homogenized in TRIzol with 3 mm tungsten carbide beads (Qiagen). Total RNA was extracted with TRIzol Reagent (Life Technologies) according to manufacturer’s instructions and quantified using the NanoDrop 1000 spectrophotometer. Retro-transcription was performed with the Reverse Transcriptase M-MLV (Life Technologies) following the manufacturer’s instructions. qRT-PCR was performed in a LightCycler 1.5 detection system (Roche) using the LightCycler FastStart DNA Master plus SYBR Green I kit (Roche) in 384-well plates according to the manufacturer’s instruction. Results are presented as normalized to the house-keeping gene (*Hprt*) and the delta-CT method was used to obtain a fold of change (FC).

The following primers were used:

*Hprt*	(F)-GTTGGATACAGGCCAGACTTTGTTG

(R)-GATTCAACTTGCGCTCATCTTAGGC

*Npy*	(F)-CCGCTCTGCGACACTACAT

(R)-TGTCTCAGGGCTGGATCTCT

*Agrp*	(F)-CGGAGGTGCTAGATCCACAGA

(R)-AGGACTCGTGCAGCCTTACAC

*Pomc*	(F)-AGTGCCAGGACCTCACCA

(R)-CAGCGAGAGGTCGAGTTTG

*Cart*	(F)-CGAGAAGAAGTACGGCCAAG

(R)-CTGGCCCCTTTCCTCACT

*Hcrt*	(F)-TTGGACCACTGCACTGAAGA

(R)-CCCAGGGAACCTTTGTAGAAG

**Quantification of eCBs by UHPLC-MS/MS**

Blood was collected before and after the last binge session and immediately centrifuged to isolate the plasma.

Brain structures (hypothalamus, VTA, dorsal striatum and nucleus accumbens) from control, bingeing and anticipating (no consumption of palatable mixture during the last day) mice were microdissected and immediately frozen in liquid nitrogen. Samples were kept at -80°C.

Samples were added to vials containing dichloromethane (8 mL), methanol (MeOH, 4 mL) (containing BHT), water (containing EDTA) and the internal standards (deuterated *N*-acylethanolamines, deuterated 2-AG), as previously described [13]. Following extraction, the lipid-containing fraction was purified by solid phase extraction (SPE). The eCBs and related NAEs were recovered from the SPE column using hexane-isopropanol 7:3 (v/v) and transferred to injection vials. The samples (1 µl) were analyzed using an Acquity UPLC® class H coupled to a Xevo TQ-S mass spectrometer (both from Waters). For the separation we used an Acquity UPLC® BEH C18 (2.1x50 mm; 1.7 µm, 40°C) column and a gradient (200 µL/min) between MeOH-H_2_O-acetic acid (75:24.9:0.1; v/v/v) and MeOH-acetic acid (99.9:0.1; v/v). Ionization was obtained using an ESI source operated in the positive mode. A quantification and a qualification transition were optimized for each analyte and MassLynx® used for data acquisition and processing. For each analyte, the ratio between the AUC of the lipid and the AUC of the corresponding internal standard was used for data normalization. Calibration curves were obtained in the same conditions and used to obtain the lipid levels. For 2-AG in brain structures lipid levels were normalized to tissues’ weight.

**Subdiaphragmatic vagotomy**

To assess the efficiency of the subdiaphragmatic vagotomy procedure (complete deafferentation) in experimental vagotmized (VGX) mice, two criteria of inclusion/exclusion were used: (*i*) a transient and reversible decrease in food intake and body weight during the first week post-vagotomy, and (*ii*) a lower anorectic response to CCK-8S (10 μg/kg, Tocris, #1166) in VGX mice compared to sham mice after an overnight fasting and a 1-hour refeeding test [14] (data not shown). Note: no differences in body weight and body composition were detected 3-4 weeks post-surgery between sham and VGX mice.

**Fiber photometry and data analysis**

The light power before entering the implanted cannula was measured with a power meter (PM100USB, Thorlabs) before the beginning of each recording session. The irradiance was ~9 mW/cm^2^. GCaMP6f- and dLight1.2-emitted fluorescent signals were collected by a femtowatt photoreceiver module (Doric Lenses). The signal was then received by the RZ5P processor (TDT). Online real-time demodulation of the fluorescence due to the 405nm and the 465 nm excitations was performed by the Synapse software (TDT). Signals were exported to Python 3.0 and analyzed off-line using TDT Python SDK packages. For the analysis, the signal processing code was inspired by previously developed pipelines [15, 16].

For the new environment paradigm, signal analysis was performed on two-time intervals: one extending from −60 to 0 seconds (home cage, HC) and the other from 0 to 60 seconds (new environment, NE).

For the tail suspension paradigm, signal analysis was performed on two-time intervals: one extending from −60 to 0 seconds (baseline) and the other from 0 to 120 seconds (tail suspension).

For GBR-evoked accumbal DA accumulation, signal analysis was performed on two-time intervals: one extending from -300 to 0 sec (baseline, before GBR administration) and the other from 0 to 1200 sec (GBR effect).

ΔF/F was calculated as [(465 nm signal_test_ − fitted 405 nm signal_ref_)/fitted 405 nm signal_ref_]. Data are presented as z-score of ΔF/F. To compare signal variations between the conditions (NE *vs* HC, tail suspension *vs* baseline, GBR *vs* baseline) for each mouse the difference between AUCs was used.

**Suppl. Fig. 1: Allostatic adaptations to time-locked palatable feeding.** (**A**) Binge consumption (ml) of palatable mixture in C57BL6 female mice during a 14-days protocol. Statistics: ***p<0.001 Binge *vs* Control, ^###^p<0.001 Binge(Last Day) *vs* Binge(First Day). (**B**) 24 hrs food intake in females considering all calories: standard diet (SD) and palatable food (PF). Statistics: ***p<0.001 Binge(SD) *vs* Control(SD), ^###^p<0.001 Binge(SD+PF) *vs* Binge(SD). (**C**) Body weight of bingeing and control female mice at the end of the time-locked palatable feeding protocol. (**D**) Longitudinal profile of fatty acid oxidation (FAO) from indirect calorimetry measurements (average of 3 consecutive days). (**E**) Real-time core temperature recording during 24 hrs and (**E^1^**) averaged values 2 hours prior and after palatable food access in control, binge, and acute animals (exposed to palatable diet for the first time). Statistics: ***p<0.001 Binge *vs* Control. (**F**) Plasma triglycerides (TG), (**G**) insulin and (**H**) corticosterone levels in animals exposed to water (Control), 1h prior (Anticipation) or 1h after (Consumption) access to palatable diet. Statistics: *p<0.05 and ***p<0.001 Anticipation *vs* Control, ^##^p<0.01 Consumption vs Anticipation. (**I, I^1^**) Blood glucose and (**J, J^1^**) insulin levels in animals daily exposed to water (Ctr) or palatable diet (binge) after an oral glucose tolerance test (OGTT). Statistics: *p<0.05 Binge *vs* Control only at 30 min post OGTT. (**K**) mRNA levels of hunger- and satiety-related genes in the hypothalamus of control and bingeing mice. For number of mice/group and statistical details see **Suppl. Table 1**.

**Suppl. Fig. 2: BE-induced molecular pathways in the dorsal striatum and nucleus accumbens.** (**A**) 1-hour consumption of water (Ctr) or palatable diet (Anticipation, Binge) during the bingeing paradigm. On day 14, “acute” animals were exposed to palatable diet for the first time while “anticipation” animals did not receive the food-reward during the last session. (**B**) Punches were extracted from the dorsal striatum (DS) and nucleus accumbens (NAc) for western blotting analysis. (**C**) Immunoblots of phosphorylated ERK1/2, ribosomal protein S6 Ser^235/236^ (P-S6^S235/236^) and S6 Ser^240/244^ (P-S6^S240/244^) in the DS and NAc. (**D**, **E**) Immunolabeling of phosphorylated S6 in the DS (**D**) and NAc (**E**) and their associated quantifications after bingeing. Scale bar: 50 μm. Statistics: ***p<0.001 Veh+Binge *vs* Veh+Control. (**F**) Immunolabeling of phosphorylated S6 in the NAc of mice pretreated (30 min) with SCH23390 or vehicle and exposed to time-locked (1h) palatable diet. Scale bar: 50 μm. For number of mice/group and statistical details see **Suppl. Table 1**.

**Suppl. Fig. 3: Effect of leptin and liraglutide on body weight and bingeing phases.** (**A, B**) Effect of leptin administration (2 injections/day for 2 consecutive days at 0.25 mg/kg) on chow food intake (**A**) and body weight (**B**). (**C**) Spontaneous locomotor activity following acute administration (i.p.) of Veh or Liraglutide (100 μg/kg) in naïve mice. (**D**) Effect of acute Veh or Liraglutide (100 μg/kg) on the locomotor activity during the anticipatory and consummatory bingeing phases. Statistics: *p<0.05, ***p<0.001 Lira+Binge (gray dots) *vs* Veh+binge (white dots). For number of mice/group and statistical details see **Suppl. Table 1**.

**Suppl. Fig. 4: Central endocannabinoids and peripheral modulation of CB1R.** Detection of 2-AG levels in the hypothalamus (**A**), ventral tegmental area (**B**), dorsal striatum (**C**) and nucleus accumbens (**D**) of control and bingeing mice (consumption and anticipatory phases). (**E**) Longitudinal measurement of energy expenditure (EE) following acute administration of AM6545 (i.p.). Note no modifications in EE. (**F**, **G**) Expression of *Cnr1* in sensory vagal neurons labeled following viral microinjections in the distal and large intestines. (**H**) cFos immunolabeling and quantification in the rostral NTS (rNTS) after acute administration of Veh or AM6545. RN indicates the reticular nucleus. Scale bar: 250 μm. For number of mice/group and statistical details see **Suppl. Table 1**.

**Suppl. Fig. 5: Homeostatic adaptations in sham and VGX mice during time-locked palatable feeding.** (**A**) 24-hours measurement of chow food intake in sham and VGX bingeing mice. Statistics: ***p<0.001 VGX+Binge *vs* Sham+Binge. (**B**) Body weight of both experimental groups. (**C-E**) Respiratory exchange ratio (RER), fatty acids oxidation (FAO) and energy expenditure (EE) in sham and VGX mice during a binge session. Statistics: *p<0.05, **p<0.01 VGX+Binge *vs* Sham+Binge. For number of mice/group and statistical details see **Suppl. Table 1**.

**Suppl. Fig. 6: Modulation of VTA DA-neurons activity.** (**A**) Effect of acute (i.p.) AM6545 or Veh on cFos-positive neurons in the DS and NAc of bingeing male (m) and female (f) mice. (**B**) Longitudinal locomotor activity induced by amphetamine (2 mg/kg, i.p.) in mice acutely (1h before) pretreated with AM6545 (AM6545+Amph) or vehicle (Veh+Amph). (**C**) Longitudinal locomotor activity induced by GBR12909 in mice acutely (1h before) pretreated with AM6545 oral gavage (po) [AM6545 (po)+GBR] or vehicle [Veh (po)+GBR]. (**D**) Immunolabeling of TH in the VTA of *Drd2*-eGFP mice. Note the co-localization between eGFP and TH in VTA DA-neurons. Scale bar: 250 μm. (**E, F**) *In vivo* recording of Ca^2+^ transients in VTA dopamine neurons of *Drd2*-Cre mice. (**E**) Ca^2+^ transients evoked following presentation of a high-fat high-sugar (HFHS) pellet (positive and reinforcing stimulus). Statistics: ***p<0.001 HFHS_after_ *vs* HFHS_before_. (**F**) Ca^2+^ transients evoked following scruff restraint (negative stimulus). Note: artefact signals while restraining the mouse were not included in the analysis. Statistics: *p<0.05 Scruff_after_ *vs* Scruff_before_. (**G**) Cumulative locomotor responses in sham and VGX mice pretreated with vehicle or AM251. Statistics: **p<0.01 Sham+AM251 *vs* Sham+Veh, and ^##^p<0.01 VGX+AM251 *vs* VGX+AM251. For number of mice/group and statistical details see **Suppl. Table 1**.
