## Supplementary material for "Identification of an endocannabinoid gut-brain vagal mechanism controlling food reward and energy homeostasis": Suppl. Table 1

**Supplementary Table 1**

| **Statistics of Figure 1** | | | | | | |
| --- | --- | --- | --- | --- | --- | --- |
| **Figure panels** | | **n** | **Statistical analysis** | | **F-value** | **p-value** |
| Fig 1B | Palatable bingeing | Ctr n=12,  Binge n=12 | 2-way ANOVA | interaction | F_(13, 308)_ = 18.68 | p < 0.0001 |
|  |  |  |  | time | F_(13, 308)_ = 19.88 | p < 0.0001 |
|  |  |  |  | groups | F_(1, 308)_ = 5755 | p < 0.0001 |
| Fig 1C | Food Intake Chow pellets | Ctr n=6,  Binge n=6 | 2-way ANOVA | interaction | F_(24, 250)_ = 2.433 | p = 0.0003 |
|  |  |  |  | time | F_(24, 250)_ = 9.989 | p < 0.0001 |
|  |  |  |  | groups | F_(1, 250)_ = 14.97 | p < 0.0001 |
| Fig 1C^1^ | Food Intake Chow pellets | Ctr n=6,  Binge n=6 | Unpaired t-test | |  | p = 0.0004,  t=5.233, df=10 |
| Fig 1D | Food Intake (all kcal)  (standard food + palatable food) | Ctr n=6,  Binge n=6 | One-way ANOVA Bonferronni *post hoc* | | F_(2, 15)_ = 23.05 | p<0.0001 |
| Fig 1E | Body weight | Ctr n=6,  Binge n=6 | 2-way ANOVA | interaction | F_(13, 130)_ = 1.544 | p = 0,1102 |
|  |  |  |  | time | F_(13, 130)_ = 3.721 | p < 0.0001 |
|  |  |  |  | groups | F_(1, 10)_ = 0.218 | p = 0.6506 |
| Fig 1E^1^ | Fat mass (%) | Ctr n=6,  Binge n=6 | Unpaired t-test | |  | p = 0.1975  t=1.38, df=10 |
| Fig 1E^1^ | Lean mass (%) | Ctr n=6,  Binge n=6 | Unpaired t-test | |  | p = 0.1047  t=1.784, df=10 |
| Fig 1F | Locomotor activity | Ctr n=6,  Binge n=6 | 2-way ANOVA | interaction | F_(96, 970)_ = 1.933 | p < 0.0001 |
|  |  |  |  | time | F_(96, 970)_ = 7.335 | p < 0.0001 |
|  |  |  |  | groups | F_(1, 970)_ = 44.83 | p < 0.0001 |
| Fig 1F^1^ | Locomotor activity | Ctr n=6,  Binge n=6 | 2-way ANOVA | interaction | F_(1, 10)_ = 8.914 | p = 0.0137 |
|  |  |  |  | time | F_(1, 10)_ = 21.37 | p = 0.0009 |
|  |  |  |  | groups | F_(1, 10)_ = 8.914 | p = 0.0061 |
| Fig 1G | RER | Ctr n=6,  Binge n=6 | 2-way ANOVA | interaction | F_(96, 970)_ = 11.38 | p < 0.0001 |
|  |  |  |  | time | F_(96, 970)_ = 32.33 | p < 0.0001 |
|  |  |  |  | groups | F_(1, 970)_ = 525.5 | p < 0.0001 |
| Fig 1G^1^ | RER | Ctr n=6,  Binge n=6 | 2-way ANOVA | interaction | F_(1, 20)_ = 1.128 | p = 0.3008 |
|  |  |  |  | time | F_(1, 20)_ = 0.0048 | p = 0.9451 |
|  |  |  |  | groups | F_(1, 20)_ = 19.53 | p = 0.0003 |
| Fig 1H | Energy Expenditure (EE) | Ctr n=6,  Binge n=6 | 2-way ANOVA | interaction | F_(96, 970)_ = 2.752 | p < 0.0001 |
|  |  |  |  | time | F_(96, 970)_ = 9.435 | p < 0.0001 |
|  |  |  |  | groups | F_(1, 970)_ = 116.9 | p < 0.0001 |
| Fig 1H^1^ | Energy Expenditure (EE) | Ctr n=6,  Binge n=6 | 2-way ANOVA | interaction | F_(1, 20)_ = 6.93 | p = 0.0160 |
|  |  |  |  | time | F_(1, 20)_ = 8.802 | p = 0.0076 |
|  |  |  |  | groups | F_(1, 20)_ = 29.94 | p < 0.0001 |
| Fig 1I | BAT Temperature | Ctr n=5,  Binge n=5 | 2-way ANOVA | interaction | F_(2, 24)_ = 0.7698 | p = 0.4742 |
|  |  |  |  | time | F_(2, 24)_ = 3.659 | p = 0.0410 |
|  |  |  |  | groups | F_(1, 24)_ = 21.24 | p = 0.0001 |
| Fig 1J | Body Core Temperature | Ctr n=8,  Binge n=8 | 2-way ANOVA | interaction | F_(96, 1344)_ = 8.638 | p < 0.0001 |
|  |  |  |  | time | F_(96, 1344)_ = 14.45 | p < 0.0001 |
|  |  |  |  | groups | F_(1, 14)_ = 9.623 | p = 0.0078 |
| Fig 1J^1^ | Body Core Temperature | Ctr n=8,  Binge n=8 | 2-way ANOVA | interaction | F_(1, 14)_ = 0.0008 | p = 0.9773 |
|  |  |  |  | time | F_(1, 14)_ = 3.492 | p = 0.0827 |
|  |  |  |  | groups | F_(1, 14)_ = 97.52 | p < 0.0001 |
| Fig 1K | Locomotor Activity | Ctr n=8,  Binge n=8 | 2-way ANOVA | interaction | F_(48, 672)_ = 8.258 | p < 0.0001 |
|  |  |  |  | time | F_(48, 672)_ = 9.059 | p < 0.0001 |
|  |  |  |  | groups | F_(1, 14)_ = 58.99 | p < 0.0001 |

| **Statistics of Figure 2** | | | | | | |
| --- | --- | --- | --- | --- | --- | --- |
| **Figure panels** | | **n** | **Statistical analysis** | | **F-value** | **p-value** |
| Fig 2A | P-ERK1/2 (DS) | Ctr n=6,  Binge n=6,  Anticip. n=6,  Acute n=6 | One-way ANOVA Bonferronni *post hoc* | | F_(3, 20)_ = 10.67 | p = 0.0002 |
|  | P-S6^S235/236^ (DS) | Ctr n=6,  Binge n=6,  Anticip. n=6,  Acute n=6 | One-way ANOVA Bonferronni *post hoc* | | F_(3, 20)_ = 4.397 | p = 0.015 |
|  | P-S6^S240/244^ (DS) | Ctr n=6,  Binge n=6,  Anticip. n=6,  Acute n=6 | One-way ANOVA Bonferronni *post hoc* | | F_(3, 20)_ = 7.935 | p = 0.0011 |
| Fig 2B | P-ERK1/2 (NAc) | Ctr n=6,  Binge n=6,  Anticip. n=6,  Acute n=6 | One-way ANOVA Bonferronni *post hoc* | | F_(3, 20)_ = 7.736 | p = 0.0013 |
|  | P-S6^S235/236^ (NAc) | Ctr n=6,  Binge n=6,  Anticip. n=6,  Acute n=6 | One-way ANOVA Bonferronni *post hoc* | | F_(3, 20)_ = 9.393 | p = 0.0004 |
|  | P-S6^S240/244^ (NAc) | Ctr n=6,  Binge n=6,  Anticip. n=6,  Acute n=6 | One-way ANOVA Bonferronni *post hoc* | | F_(3, 20)_ = 6.546 | p = 0.0029 |
| Fig 2C | GBR-induced locomotor activity | Ctr+GBR n=5,  NoBinge+GBR n=5 | 2-way ANOVA | interaction | F_(23, 96)_ = 1.565 | p = 0.0686 |
|  |  |  |  | time | F_(23, 96)_ = 9.368 | p < 0.0001 |
|  |  |  |  | groups | F_(1, 96)_ = 2.552 | p = 0.1134 |
| Fig 2C^1^ | GBR-induced locomotor activity | Ctr+GBR n=5,  NoBinge+GBR n=5 | Unpaired t-test | |  | p = 0.4505  t=0.7934, df=8 |
| Fig 2D | GBR-induced locomotor activity post binge-exposure | Ctr+GBR n=5,  Binge+GBR n=5 | 2-way ANOVA | interaction | F_(23, 96)_ = 10.17 | p < 0.0001 |
|  |  |  |  | time | F_(23, 96)_ = 12.08 | p < 0.0001 |
|  |  |  |  | groups | F_(1, 96)_ = 49.5 | p < 0.0001 |
| Fig 2D^1^ | GBR-induced locomotor activity post binge-exposure | Ctr+GBR n=5,  Binge+GBR n=5 | Unpaired t-test | |  | p = 0.0082,  t=3.49, df=8 |
| Fig 2E | Effect of SCH on BE | Ctr n=4,  Veh+Binge n=5,  SCH+Binge n=5 | 2-way ANOVA | interaction | F_(4, 33)_ = 54.53 | p < 0.0001 |
|  |  |  |  | time | F_(2, 33)_ = 108.8 | p < 0.0001 |
|  |  |  |  | groups | F_(2, 33)_ = 245.9 | p < 0.0001 |
| Fig 2F | Effect of Halo on BE | Ctr n=8,  Halo0.25 n=6,  Halo0.5 n=6 | One-way ANOVA Bonferronni *post hoc* | | F_(2, 17)_ = 1.01 | p = 0.3850 |
| Fig 2G^1^ | P-S6^S235/236^ (DS) | Ctr n=5,  Veh+Binge n=6,  SCH+Binge n=6 | One-way ANOVA Bonferronni *post hoc* | | F_(2, 14)_ = 53.06 | p < 0.0001 |
| Fig 2H^1^ | P-S6^S240/244^ (DS) | Ctr n=5,  Veh+Binge n=6,  SCH+Binge n=6 | One-way ANOVA Bonferronni *post hoc* | | F_(2, 14)_ = 92.71 | p < 0.0001 |
| Fig 2G^2^ | P-S6^S235/236^ (NAc) | Ctr n=5,  Veh+Binge n=6,  SCH+Binge n=6 | One-way ANOVA Bonferronni *post hoc* | | F_(2, 14)_ = 89.16 | p < 0.0001 |
| Fig 2H^2^ | P-S6^S240/244^ (NAc) | Ctr n=5,  Veh+Binge n=6,  SCH+Binge n=6 | One-way ANOVA Bonferronni *post hoc* | | F_(2, 14)_ = 110.6 | p < 0.0001 |
| Fig 2I | Locomotor activity | Veh+Binge n=6,  SCH+Binge n=6 | 2-way ANOVA | interaction | F_(48, 480)_ = 1.156 | p = 0.2275 |
|  |  |  |  | time | F_(48, 480)_ = 4.138 | p < 0.0001 |
|  |  |  |  | groups | F_(1, 10)_ = 1.98 | p = 0.1897 |
| Fig 2I^1^ | Locomotor activity | Veh+Binge n=6,  SCH+Binge n=6 | 2-way ANOVA | interaction | F_(1, 10)_ = 12.07 | p = 0.0060 |
|  |  |  |  | time | F _1, 10)_ = 9.891 | p = 0.0104 |
|  |  |  |  | groups | F_(1, 10)_ = 8.607 | p = 0.0149 |
| Fig 2J | Food intake | Veh+Binge n=6,  SCH+Binge n=6 | 2-way ANOVA | interaction | F_(96, 960)_ = 3.878 | p < 0.0001 |
|  |  |  |  | time | F_(96, 960)_ = 201 | p < 0.0001 |
|  |  |  |  | groups | F_(1, 10)_ = 2.438 | p = 0.046 |
| Fig 2K | Locomotor activity | Ctr+SKF n=6,  Binge+SKF n=6 | 2-way ANOVA | interaction | F_(23, 230)_ = 3.031 | p < 0.0001 |
|  |  |  |  | time | F_(23, 230)_ = 21.59 | p < 0.0001 |
|  |  |  |  | groups | F_(1, 10)_ = 12.05 | p = 0.0006 |
| Fig 2K^1^ | Locomotor activity (2 hrs) | Ctr+SKF n=6,  Binge+SKF n=6 | Unpaired t-test | |  | p = 0.5415  t=0.6321, df=10 |
|  | Locomotor activity (30 min) | Ctr+SKF n=6,  Binge+SKF n=6 | Unpaired t-test | |  | p = 0.0135  t=2.991, df=10 |

| **Statistics of Figure 3** | | | | | | |
| --- | --- | --- | --- | --- | --- | --- |
| **Figure panels** | | **n** | **Statistical analysis** | | **F-value** | **p-value** |
| Fig 3A | Palatable bingeing | Veh n=12,  Leptin n=6,  Insulin n=6  Exe-4 n=6  Lira n=6  CCK-8S n=6  AM251 n=6 | One-way ANOVA Bonferronni *post hoc* | | F_(6, 41)_ = 47.38 | p < 0.0001 |
| Fig 3B | AEA | before=6,  after n=6 | Unpaired t-test | |  | p = 0.8585  t=0.183, df=10 |
|  | 2-AG | before=6,  after n=6 | Unpaired t-test | |  | p < 0.0001  t=8.161, df=10 |
|  | DHEA | before n=6,  after n=6 | Unpaired t-test | |  | p = 0.7251  t=0.3617, df=10 |
|  | OEA | before=6,  after n=6 | Unpaired t-test | |  | p = 0.5115  t=0.6808, df=10 |
| Fig  3C | Palatable bingeing | Veh+Binge  n=6,  AM6545+  Binge n=6,  JD5037+  Binge n=6 | One-way ANOVA Bonferronni *post hoc* | | F_(2, 15)_ = 110.06 | p < 0.0001 |
| Fig 3D | Palatable bingeing | All Binge n=20,  Veh n=5,  AM6545 n=5,  JZL184 n=5,  AM+JZL n=5 | 2-way ANOVA | interaction | F_(3, 16)_ = 167.8 | p < 0.0001 |
|  |  |  |  | time | F_(1, 16)_ = 168.7 | p < 0.0001 |
|  |  |  |  | groups | F_(3, 16)_ = 57.9 | p < 0.0001 |
| Fig 3E | Palatable bingeing | Veh+Binge n=5,  AM+Binge n=5,  JZL+Binge n=5 | 2-way ANOVA | interaction | F_(8, 60)_ = 22.54 | p < 0.0001 |
|  |  |  |  | time | F_(4, 60)_ = 5.554 | p = 0.0007 |
|  |  |  |  | groups | F_(2, 60)_ = 337.9 | p < 0.0001 |
| Fig 3F | Body Core Temperature | Veh+Binge n=8,  AM+Binge n=8 | 2-way ANOVA | interaction | F_(48, 672)_ = 5.557 | p < 0.0001 |
|  |  |  |  | time | F_(48, 672)_ = 9.794 | p < 0.0001 |
|  |  |  |  | groups | F_(1, 14)_ = 4.839 | p = 0.0451 |
| Fig 3G | Locomotor Activity | Veh+Binge n=8,  AM+Binge n=8 | 2-way ANOVA | interaction | F_(48, 672)_ = 2,641 | p < 0.0001 |
|  |  |  |  | time | F_(48, 672)_ = 9.043 | p < 0.0001 |
|  |  |  |  | groups | F_(1, 14)_ = 6.052 | p = 0.0275 |
| Fig 3H | Fatty acid oxidation (FAO) | Veh n=6,  AM6545 n=5 | 2-way ANOVA | interaction | F_(192, 1728)_ = 5.28 | p < 0.0001 |
|  |  |  |  | time | F_(192, 1728)_ = 19.2 | p < 0.0001 |
|  |  |  |  | groups | F_(1, 9)_ = 17.5 | p = 0.0024 |
| Fig 3H^1^ | Fatty acid oxidation (FAO) | Veh n=6,  AM6545 n=5 | 2-way ANOVA | interaction | F_(1, 18)_ = 7.789 | p = 0.0121 |
|  |  |  |  | time | F_(1, 18)_ = 8.961 | p = 0.0078 |
|  |  |  |  | groups | F_(1, 18)_ = 84.11 | p < 0.0001 |
| Fig 3H^2^ | Ratio FAO/Food Intake | Veh n=6,  AM6545 n=5 | 2-way ANOVA | interaction | F_(1, 18)_ = 0.057 | p = 0.8125 |
|  |  |  |  | time | F_(1, 18)_ = 0.705 | p = 0.4119 |
|  |  |  |  | groups | F_(1, 18)_ = 99.82 | p < 0.0001 |
| Fig 3I | Palatable bingeing | Veh (po) n=5,  AM6545 (po) n=5 | Unpaired t-test | |  | p = 0.4263,  t=0.8847, df=4 |
| Fig 3J | Fatty acid oxidation (FAO) | Veh n=6,  AM6545 n=6 | 2-way ANOVA | interaction | F_(95, 950)_ = 0.479 | p > 0.9999 |
|  |  |  |  | time | F_(95, 950)_ = 15.77 | p < 0.0001 |
|  |  |  |  | groups | F_(1, 10)_ = 0.134 | p = 0.7219 |
| Fig 3K | cFos (NTS) | Veh+Ctr  n=6,  Veh+Binge  n=6,  AM6545  +Binge n=6 | One-way ANOVA Bonferronni *post hoc* | | F_(2, 15)_ = 49.74 | p < 0.0001 |
|  | cFos (AP) | Veh+Ctr  n=6,  Veh+Binge  n=6,  AM6545  +Binge n=6 | One-way ANOVA Bonferronni *post hoc* | | F_(2, 15)_ = 99.63 | p < 0.0001 |
| Fig 3L | cFos (lPBN) | Veh+Ctr  n=6,  Veh+Binge  n=6,  AM6545  +Binge n=6 | One-way ANOVA Bonferronni *post hoc* | | F_(2, 15)_ = 106.3 | p < 0.0001 |

| **Statistics of Figure 4** | | | | | | | | |
| --- | --- | --- | --- | --- | --- | --- | --- | --- |
| **Figure panels** | | **n** | **Statistical analysis** | | | | **F-value** | **p-value** |
| Fig 4C | Palatable bingeing  (Sham *vs* VGX, JZL184) | Sham  +JZL184 n=6,  VGX  +JZL184  n=6 | 2-way ANOVA | | interaction | | F_(4, 40)_ = 5.543 | p = 0.0012 |
|  |  |  |  |  | time | | F_(4, 40)_ = 12.01 | p < 0.0001 |
|  |  |  |  |  | groups | | F_(1, 10)_ = 55.27 | p < 0.0001 |
| Fig 4D^2^ | AP cFos | Sham  +AM6545 n=5,  VGX  +AM6545 n=5 | Unpaired t-test | | | |  | p < 0.0001  t=9.945, df=8 |
|  | NTS cFos |  |  |  |  |  |  | p < 0.0001  t=7.777, df=8 |
|  | lPBN cFos |  |  |  |  |  |  | p < 0.0001  t=8.337, df=8 |
| Fig 4E | Palatable bingeing | Sham+Veh n=6,  Sham  +AM6545 n=6,  VGX+Veh n=5,  VGX  +AM6545 n=5 | 2-way ANOVA | interaction | | | F_(1, 9)_ = 107.3 | p < 0.0001 |
|  |  |  |  | time | | | F_(1, 9)_ = 156.7 | p < 0.0001 |
|  |  |  |  | groups | | | F_(1, 9)_ = 4.271 | p = 0.0687 |
| Fig 4F | Fatty acid oxidation (FAO) | Sham+Veh n=6,  Sham  +AM6545 n=6 | 2-way ANOVA | interaction | | | F_(96, 960)_ = 4.425 | p < 0.0001 |
|  |  |  |  | time | | | F_(96, 960)_ = 13.08 | p < 0.0001 |
|  |  |  |  | groups | | | F_(1, 10)_ = 10.13 | p = 0.0098 |
| Fig 4F^1^ | Ratio FAO/Food Intake | Sham+Veh n=6,  Sham  +AM6545 n=6 | Unpaired t-test | | | |  | p < 0.0001  t=4.626, df=10 |
| Fig 4G | Fatty acid oxidation (FAO) | VGX+Veh n=6,  VGX  +AM6545 n=6 | 2-way ANOVA | | | interaction | F_(96, 960)_ = 1.488 | p = 0.0025 |
|  |  |  |  |  |  | time | F_(96, 960)_ = 12.1 | p < 0.0001 |
|  |  |  |  |  |  | groups | F_(1, 10)_ = 3.709 | p = 0.0830 |
| Fig 4G^1^ | Ratio FAO/Food Intake | VGX+Veh n=6,  VGX+AM6545 n=6 | Unpaired t-test | | | |  | p = 0.2838  t=1.133, df=10 |
| Fig 4H | cFos in PVN | Veh n=5,  Sham+AM6545 n=5,  VGX+AM6545 n=5 | One-way ANOVA Bonferronni *post hoc* | | | | F_(2, 12)_ = 155.8 | p < 0.0001 |
| Fig 4I | cFos in DMH | Veh n=5,  Sham+AM6545 n=5,  VGX+AM6545 n=5 | One-way ANOVA Bonferronni *post hoc* | | | | F_(2, 12)_ = 63.71 | p < 0.0001 |

| **Statistics of Figure 5** | | | | | | |
| --- | --- | --- | --- | --- | --- | --- |
| **Figure panels** | | **n** | **Statistical analysis** | | **F-value** | **p-value** |
| Fig  5A | P-S6^S235/236^ (DS) | Veh+Ctr  n=6,  Veh+Binge  n=6,  AM6545  +Binge n=6 | One-way ANOVA Bonferronni *post hoc* | | F_(2, 15)_ = 176 | p < 0.0001 |
|  | P-S6^S235/236^ (NAc) | Veh+Ctr  n=6,  Veh+Binge  n=6,  AM6545  +Binge n=6 | One-way ANOVA Bonferronni *post hoc* | | F_(2, 15)_ = 49.73 | p < 0.0001 |
|  | cFos (DS) | Veh+Ctr  n=6,  Veh+Binge  n=6,  AM6545  +Binge n=6 | One-way ANOVA Bonferronni *post hoc* | | F_(2, 15)_ = 159.2 | p < 0.0001 |
|  | cFos (NAc) | Veh+Ctr  n=6,  Veh+Binge  n=6,  AM6545  +Binge n=6 | One-way ANOVA Bonferronni *post hoc* | | F_(2, 15)_ = 132.9 | p < 0.0001 |
| Fig 5B | GBR-induced locomotor activity (AM6545) | Veh+GBR n=5,  AM6545  +GBR n=5 | 2-way ANOVA | interaction | F_(23, 184)_ = 6.443 | p < 0.0001 |
|  |  |  |  | time | F_(23, 184)_ = 9.964 | p < 0.0001 |
|  |  |  |  | groups | F_(1, 8)_ = 9.723 | p = 0.0143 |
| Fig 5B^1^ | GBR-induced locomotor activity (AM6545) | Veh+GBR n=5,  AM6545  +GBR n=5 | Unpaired t-test | |  | p = 0.0147  t=3.096, df=8 |
| Fig 5C | GBR-induced locomotor activity (JD5037) | Veh+GBR  n=7,  JD5037  +GBR n=7 | 2-way ANOVA | interaction | F_(17, 204)_ = 6.485 | p < 0.0001 |
|  |  |  |  | time | F_(17, 204)_ = 13.71 | p < 0.0001 |
|  |  |  |  | groups | F_(1, 12)_ = 8.566 | p = 0.0127 |
| Fig 5C^1^ | GBR-induced locomotor activity (JD5037) | Veh+GBR n=5,  AM6545  +GBR n=5 | Unpaired t-test | |  | p = 0.0111  t=2.998, df=12 |
| Fig 5D | cFos | Veh+GBR n=5,  AM6545  +GBR n=5 | Unpaired t-test | |  | p < 0.0001  t=5.615, df=8 |
| Fig 5E^1^ | Dopamine dynamics  (z-Score, LIght1.2) | Veh+GBR n=6,  AM6545  +GBR n=6 | 2-way ANOVA | interaction | F_(4, 40)_ = 11.22 | p < 0.0001 |
|  |  |  |  | time | F_(4, 40)_ = 7.495 | p = 0.0001 |
|  |  |  |  | groups | F_(1, 10)_ = 3.439 | p = 0.0025 |
| Fig 5F | Amph-induced locomotor activity (AM6545) | Veh  +Amph n=6,  AM6545  +Amph n=6 | Unpaired t-test | |  | p = 0.9876  t=0.0159, df=10 |
| Fig 5G | Amph-induced locomotor activity (AM251) | Veh+  Amph n=6,  AM251+  Amph n=6 | Unpaired t-test | |  | p = 0.0003  t=5.417, df=10 |
| Fig 5H | GBR-induced locomotor activity (VGX animals) | VGX+GBR n=9,  VGX+AM+GBR n=9 | 2-way ANOVA | interaction | F_(17, 144)_ = 0,465 | p = 0.9644 |
|  |  |  |  | time | F_(17, 144)_ = 13,69 | p < 0.0001 |
|  |  |  |  | groups | F_(1, 144)_ = 0,299 | p = 0.5853 |
| Fig 5H^1^ | GBR-induced locomotor activity | VGX+GBR n=9,  VGX+AM+GBR n=9 | Unpaired t-test | |  | p = 0.7320  t=0.3485, df=16 |
| Fig 5I | GBR-induced locomotor activity (po) | Veh+GBR n=6,  AM6545+GBR n=6 | Unpaired t-test | |  | p = 0.8694  t=0.1687, df=10 |

| **Statistics of Figure 6** | | | | | | |
| --- | --- | --- | --- | --- | --- | --- |
| **Figure panels** | | **n** | **Statistical analysis** | | **F-value** | **p-value** |
| Fig 6C | z-Score Ca^2+^  (Tail suspension) | Veh n=5,  AM6545 n=5 | Paired t-test | |  | p = 0.0038 t=6.037, df=4 |
| Fig 6D | z-Score Ca^2+^  (New environment) | Veh n=5,  AM6545 n=5 | Paired t-test | |  | p = 0.0479  t=2.818, df=4 |
| Fig 6E | Time of immobility | Sham+Veh n=7,  Sham  +AM6545 n=7,  VGX+Veh n=7,  VGX  +AM6545 n=7 | 2-way ANOVA | interaction | F_(1, 12)_ = 3.451 | p = 0.0879 |
|  |  |  |  | treatment | F_(1, 12)_ = 0.5823 | p = 0.4602 |
|  |  |  |  | groups | F_(1, 12)_ = 0.5631 | p = 0.4675 |
| Fig 6F | Locomotor activity  (AM6545, new cage) | Sham+Veh n=7,  Sham  +AM6545 n=7,  VGX+Veh n=7,  VGX  +AM6545 n=7 | 2-way ANOVA | interaction | F_(87, 696)_ = 2.045 | p < 0.0001 |
|  |  |  |  | time | F_(29, 696)_ = 8.599 | p < 0.0001 |
|  |  |  |  | groups | F_(3, 24)_ = 8.132 | p = 0.0007 |
| Fig 6F^1^ | Locomotor activity  (AM6545, new cage) | Sham+Veh n=7,  Sham  +AM6545 n=7,  VGX+Veh n=7,  VGX  +AM6545 n=7 | 2-way ANOVA | interaction | F_(1, 12)_ = 5.775 | p = 0.0333 |
|  |  |  |  | treatment | F_(1,12)_ = 12.51 | p = 0.0041 |
|  |  |  |  | groups | F_(1, 12)_ = 4.895 | p = 0.0471 |

| **Statistics of Suppl. Figure 1** | | | | | | |
| --- | --- | --- | --- | --- | --- | --- |
| **Figure panels** | | **n** | **Statistical analysis** | | **F-value** | **p-value** |
| SFig 1A | Palatable bingeing in female mice | Ctr n=6,  Binge n=6 | 2-way ANOVA | interaction | F_(1, 10)_ = 51.73 | p < 0.0001 |
|  |  |  |  | time | F_(1, 10)_ = 54.22 | p < 0.0001 |
|  |  |  |  | groups | F_(1, 10)_ = 343.6 | p < 0.0001 |
| SFig 1B | Food Intake (all kcal)  (standard food + palatable food) female mice | Ctr n=6,  Binge n=6 | One-way ANOVA Bonferronni *post hoc* | | F_(2, 15)_ = 19.91 | p<0.0001 |
| SFig 1C | Body weight (female mice) | Ctr n=6,  Binge n=6 | Unpaired t-test | |  | p = 0.5701  t=0.5872, df=10 |
| SFig 1D | Fatty acid oxidation (FAO) | Ctr n=6,  Binge n=6 | 2-way ANOVA | interaction | F_(96, 960)_ = 10.66 | p < 0.0001 |
|  |  |  |  | time | F_(96, 960)_ = 29.67 | p < 0.0001 |
|  |  |  |  | groups | F_(1, 10)_ = 152.3 | p < 0.0001 |
| SFig 1E | Core Body Temperature | Ctr n=8,  Binge n=8,  Acute n=8 | 2-way ANOVA | interaction | F_(192, 1358)_ = 6,36 | p < 0.0001 |
|  |  |  |  | time | F_(96, 679)_ = 16,57 | p < 0.0001 |
|  |  |  |  | groups | F_(2, 1358)_ = 33,24 | p < 0.0001 |
| SFig 1E^1^ | Core Body Temperature | Ctr n=8,  Binge n=8,  Acute n=8 | 2-way ANOVA | interaction | F_(2, 21)_ = 48.57 | p < 0.0001 |
|  |  |  |  | time | F_(1, 21)_ = 22.8 | p < 0.0001 |
|  |  |  |  | groups | F_(2, 21)_ = 74.9 | p < 0.0001 |
| SFig 1F | Plasma TG | Ctr n=5,  Antic. n=6,  Cons. n=6 | One-way ANOVA Bonferronni *post hoc* | | F_(2, 14)_ = 7.56 | p = 0.0059 |
| SFig 1G | Plasma Insulin | Ctr n=5,  Antic. n=6,  Cons. n=6 | One-way ANOVA Bonferronni *post hoc* | | F_(2, 14)_ = 6.327 | p = 0.0110 |
| SFig 1H | Plasma Corticosterone | Ctr n=5,  Antic. n=6,  Cons. n=6 | One-way ANOVA Bonferronni *post hoc* | | F_(2, 14)_ = 10.65 | p = 0.0015 |
| SFig 1I | Glucose (OGTT) | Ctr n=12,  Binge n=12 | 2-way ANOVA | interaction | F_(6, 132)_ = 3.781 | p = 0.0017 |
|  |  |  |  | time | F_(6, 132)_ = 91.12 | p < 0.0001 |
|  |  |  |  | groups | F_(1, 22)_ = 0.0018 | p = 0.9662 |
| SFig 1I^1^ | Glucose (AUC) | Ctr n=12,  Binge n=12 | Unpaired t-test | |  | p = 0.3629,  t=0.9291, df=22 |
| SFig 1J | Insulin (OGTT) | Ctr n=12,  Binge n=12 | 2-way ANOVA | interaction | F_(3, 66)_ = 0.8114 | p = 0.4921 |
|  |  |  |  | time | F_(3, 66)_ = 42.81 | p < 0.0001 |
|  |  |  |  | groups | F_(1, 22)_ = 2.638 | p = 0.1186 |
| SFig 1J^1^ | Insulin (AUC) | Ctr n=12,  Binge n=12 | Unpaired t-test | |  | p = 0.1455,  t=1.509, df=22 |
| SFig 1K | *Npy* | Ctr n=5,  Binge n=6 | Unpaired t-test | |  | p = 0.6653,  t=0.4471, df=9 |
|  | *Agrp* | Ctr n=5,  Binge n=6 | Unpaired t-test | |  | p = 0.1706,  t=1.489, df=9 |
|  | *Pomc* | Ctr n=5,  Binge n=6 | Unpaired t-test | |  | p = 0.9383,  t=0.0796, df=9 |
|  | *Cart* | Ctr n=5,  Binge n=6 | Unpaired t-test | |  | p = 0.6415,  t=0.4817, df=9 |
|  | *Hcrt* | Ctr n=5,  Binge n=6 | Unpaired t-test | |  | p = 0.4290,  t=0.8281, df=9 |

| **Statistics of Suppl. Figure 2** | | | | | |
| --- | --- | --- | --- | --- | --- |
| **Figure panels** | | **n** | **Statistical analysis** | **F-value** | **p-value** |
| SFig 2D | P-S6^S235/236^-cells (DS) | Ctr n=4,  Binge n=5 | Unpaired t-test |  | p = 0.0002,  t=7.261, df=7 |
| SFig 2E | P-S6^S235/236^-cells (NAc) | Ctr n=4,  Binge n=5 | Unpaired t-test |  | p = 0.0002,  t=7.385, df=7 |

| **Statistics of Suppl. Figure 3** | | | | | | |
| --- | --- | --- | --- | --- | --- | --- |
| **Figure panels** | | **n** | **Statistical analysis** | | **F-value** | **p-value** |
| SFig 3A | Food intake (leptin) | Ctr n=6,  Binge n=6  before *vs* after treatment | Paired t-test | |  | p = 0.0017,  t=6.093, df=5 |
| SFig 3B | Body weight (leptin) | Ctr n=6,  Binge n=6  before *vs* after treatment | Paired t-test | |  | p = 0.0013,  t=6.445, df=5 |
| SFig 3C | Spontaneous locomotor activity (liraglutide) | Veh n=6  Lira n=6 | 2-way ANOVA | interaction | F_(24, 240)_ = 0.995 | p = 0.4731 |
|  |  |  |  | time | F_(24, 240)_ = 4.004 | p < 0.0001 |
|  |  |  |  | groups | F_(1, 10)_ = 1.243 | p = 0.2643 |
| SFig 3D | BE-induced locomotor activity (liraglutide) | Veh n=6  Lira n=6 | 2-way ANOVA | interaction | F_(24, 240)_ = 2.531 | p = 0.0002 |
|  |  |  |  | time | F_(24, 240)_ = 11.72 | p < 0.0001 |
|  |  |  |  | groups | F_(1, 10)_ = 9.105 | p = 0.0130 |

| **Statistics of Suppl. Figure 4** | | | | | | |
| --- | --- | --- | --- | --- | --- | --- |
| **Figure panels** | | **n** | **Statistical analysis** | | **F-value** | **p-value** |
| SFig 4A | 2-AG (Hypothalamus) | Ctr n=7,  Binge n=7,  Anticip. n=6 | One-way ANOVA Bonferronni *post hoc* | | F_(2, 17)_ = 0.6732 | p = 0.5231 |
| SFig 4B | 2-AG (VTA) | Ctr n=7,  Binge n=7,  Anticip. n=6 | One-way ANOVA Bonferronni *post hoc* | | F_(2, 17)_ = 0.3586 | p = 0.7038 |
| SFig 4C | 2-AG (DS) | Ctr n=7,  Binge n=7,  Anticip. n=6 | One-way ANOVA Bonferronni *post hoc* | | F_(2, 17)_ = 0.477 | p = 0.6287 |
| SFig 4D | 2-AG (NAc) | Ctr n=7,  Binge n=7,  Anticip. n=6 | One-way ANOVA Bonferronni *post hoc* | | F_(2, 17)_ = 0.5005 | p = 0.6149 |
| SFig 4E | Energy Expenditure  AM6545 | Veh n=6,  AM6545 n=5 | 2-way ANOVA | interaction | F_(96, 864)_ = 1,236 | p = 0.0703 |
|  |  |  |  | time | F_(96, 864)_ = 4,304 | p < 0.0001 |
|  |  |  |  | groups | F_(1, 9)_ = 1,835 | p = 0.2085 |
| SFig 4H | cFos in the rostral NTS | Veh n=6,  AM6545 n=6 | Unpaired t-test | |  | p = 0.8000,  t=0.2602, df=10 |

| **Statistics of Suppl. Figure 5** | | | | | | |
| --- | --- | --- | --- | --- | --- | --- |
| **Figure panels** | | **n** | **Statistical analysis** | | **F-value** | **p-value** |
| SFig 5A | Food Intake | Sham/Binge n=6,  VGX/Binge n=5 | 2-way ANOVA | interaction | F_(96, 864)_ = 15.2 | p < 0.0001 |
|  |  |  |  | time | F_(96, 864)_ = 348.1 | p < 0.0001 |
|  |  |  |  | groups | F_(1, 9)_ = 8.109 | p = 0.0192 |
| SFig 5B | Body weight | Sham/Binge n=6,  VGX/Binge n=5 | Unpaired t-test | |  | p = 0.5254  t=0.6606, df=9 |
| SFig 5C | RER | Sham/Binge n=6,  VGX/Binge n=5 | 2-way ANOVA | interaction | F_(1, 9)_ = 12.42 | p = 0.0065 |
|  |  |  |  | time | F_(1, 9)_ = 62.5 | p < 0.0001 |
|  |  |  |  | groups | F_(1, 9)_ = 2.472 | p = 0.1504 |
| SFig 5D | FAO | Sham/Binge n=6,  VGX/Binge n=5 | 2-way ANOVA | interaction | F_(1, 9)_ = 15.02 | p = 0.0038 |
|  |  |  |  | time | F_(1, 9)_ = 46.32 | p < 0.0001 |
|  |  |  |  | groups | F_(1, 9)_ = 3.213 | p = 0.1066 |
| SFig 5E | EE | Sham/Binge n=6,  VGX/Binge n=5 | 2-way ANOVA | interaction | F_(1, 9)_ = 4.006 | p = 0.0764 |
|  |  |  |  | time | F_(1, 9)_ = 99.22 | p < 0.0001 |
|  |  |  |  | groups | F_(1, 9)_ = 0.359 | p = 0.5638 |

| **Statistics of Suppl. Figure 6** | | | | | | |
| --- | --- | --- | --- | --- | --- | --- |
| **Figure panels** | | **n** | **Statistical analysis** | | **F-value** | **p-value** |
| SFig 6A | cFos in the DS  (males *vs* females) | Ctr n=6  (3m & 3f)  Binge n=6  (3m & 3f)  AM6545  +Binge n=6  (3m & 3f) | 2-way ANOVA | interaction | F_(2, 12)_ = 1.037 | p = 0.3843 |
|  |  |  |  | sex | F_(1, 12)_ = 0.2177 | p = 0.6492 |
|  |  |  |  | groups | F_(2, 12)_ = 151.6 | p < 0.0001 |
|  | cFos in the NAc  (males *vs* females) | Ctr n=6  (3m & 3f)  Binge n=6  (3m & 3f)  AM6545  +Binge n=6  (3m & 3f) | 2-way ANOVA | interaction | F_(2, 12)_ = 0.7995 | p = 0.4721 |
|  |  |  |  | sex | F_(1, 12)_ = 0.7196 | p = 0.4129 |
|  |  |  |  | groups | F_(2, 12)_ = 126.9 | p < 0.0001 |
| SFig 6B | Amph-induced locomotor activity | Veh+Amph n=6,  AM6545  +Amph n=6 | 2-way ANOVA | interaction | F_(17, 90)_ = 0,7921 | p = 0.6976 |
|  |  |  |  | time | F_(17, 90)_ = 30,62 | p < 0.0001 |
|  |  |  |  | groups | F_(1, 90)_ = 0,0106 | p = 0.9180 |
| SFig 6C | GBR-induced locomotor activity (po) | Veh+GBR n=6,  AM6545  +GBR n=6 | 2-way ANOVA | interaction | F_(17, 90)_ = 0,7426 | p = 0.7513 |
|  |  |  |  | time | F_(17, 90)_ = 3,449 | p < 0.0001 |
|  |  |  |  | groups | F_(1, 90)_ = 2,981 | p = 0.0877 |
| SFig 6E | HFHS (positive valence) | n=5 | Paired t-test | |  | p = 0.0005  t=10.37, df=4 |
| SFig 6F | Scruff restraint  (negative valence) | n=5 | Paired t-test | |  | p = 0.0222  t=3.63, df=4 |
| SFig 6G | Locomotor activity  (AM251, new environment) | Sham+Veh n=7,  Sham  +AM6545 n=7,  VGX+Veh n=7,  VGX  +AM6545 n=7 | 2-way ANOVA | interaction | F_(1, 12)_ = 0.0698 | p = 0.7961 |
|  |  |  |  | treatment | F_(1,12)_ = 31.44 | p = 0.0001 |
|  |  |  |  | groups | F_(1, 12)_ = 0.0079 | p = 0.9303 |
