## Supplementary figures and images for "Identification of an endocannabinoid gut-brain vagal mechanism controlling food reward and energy homeostasis"

### Suppl. Figure 1

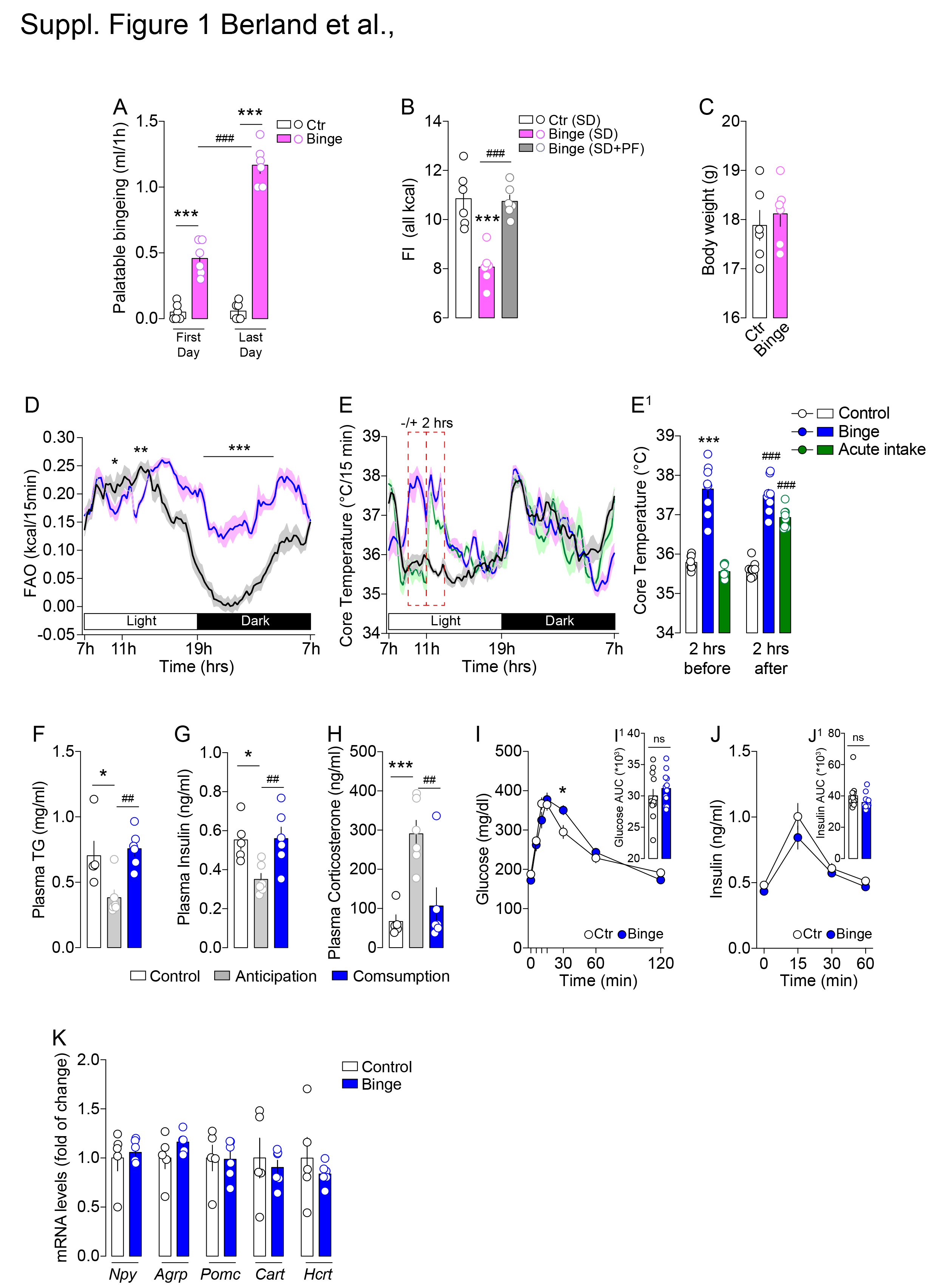

### Suppl. Figure 2

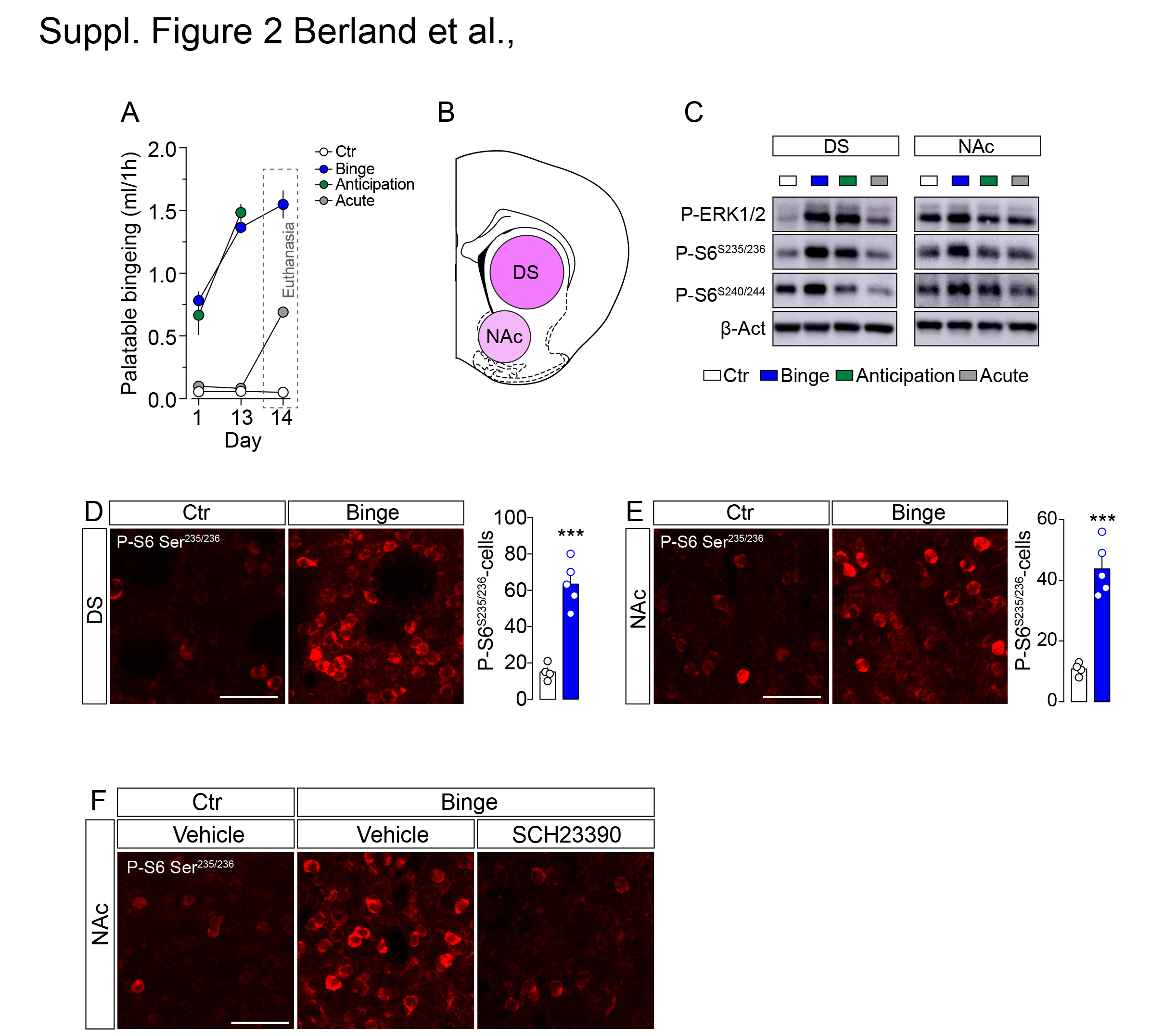

### Suppl. Figure 3

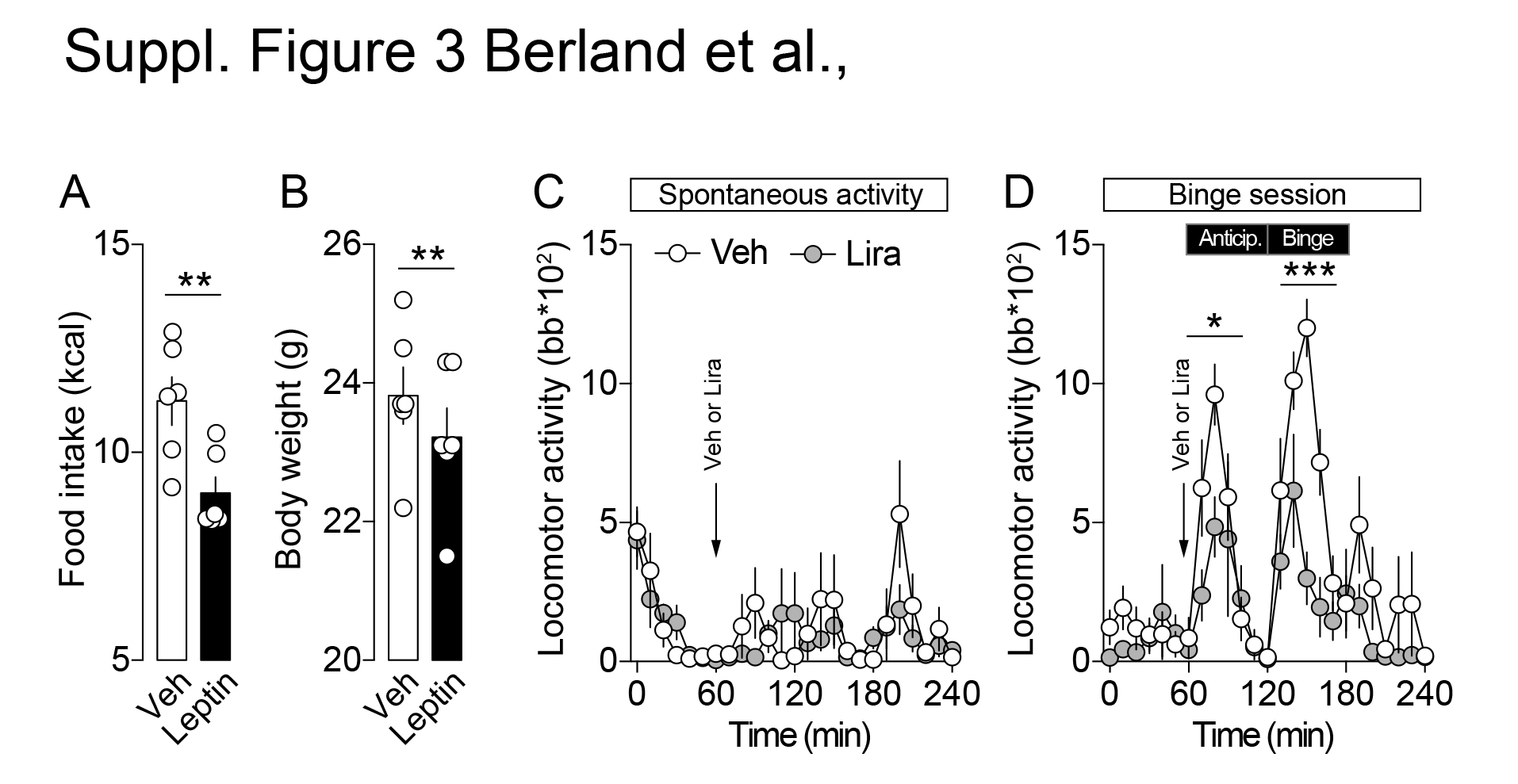

### Suppl. Figure 4

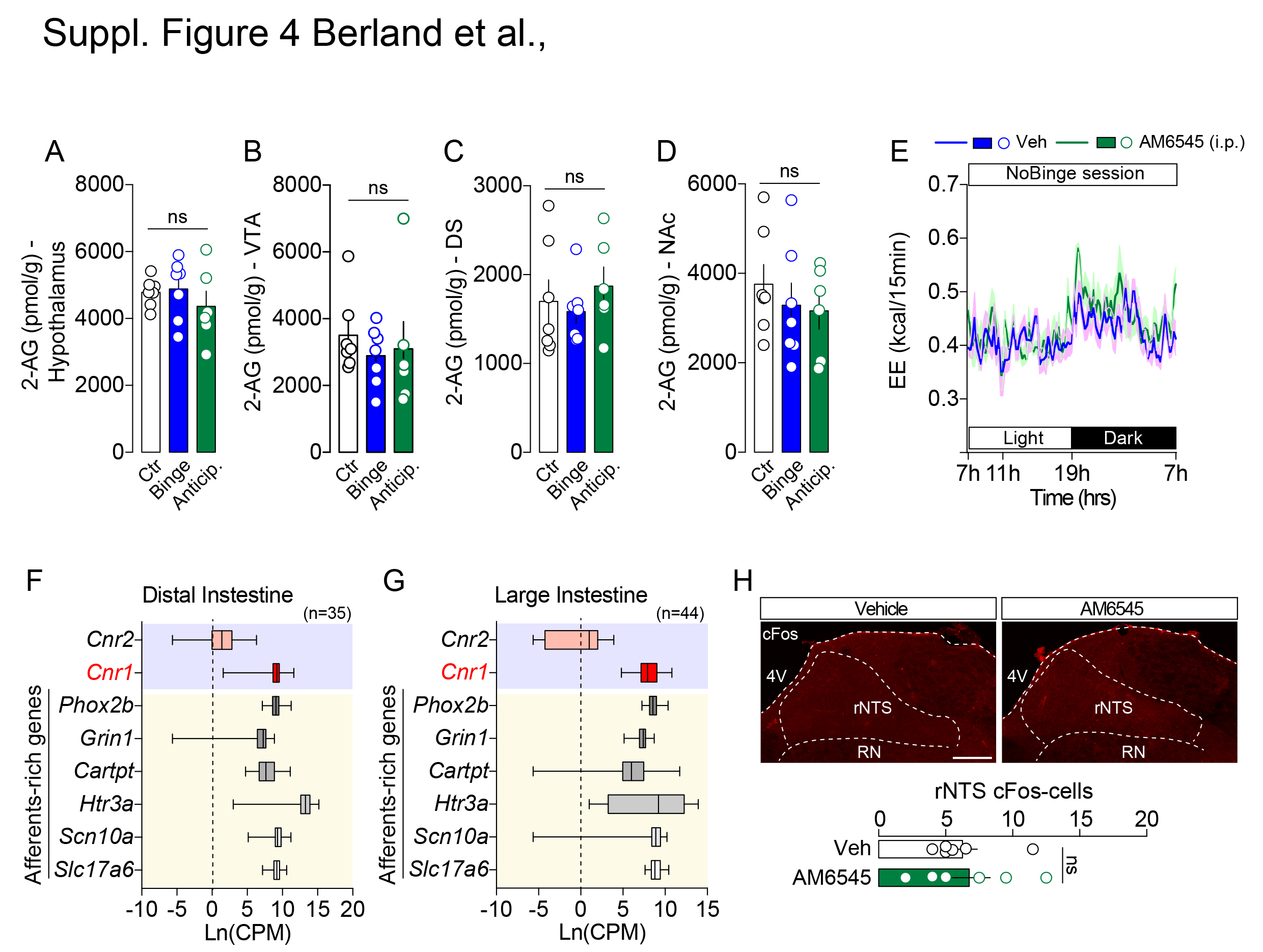

### Suppl. Figure 5

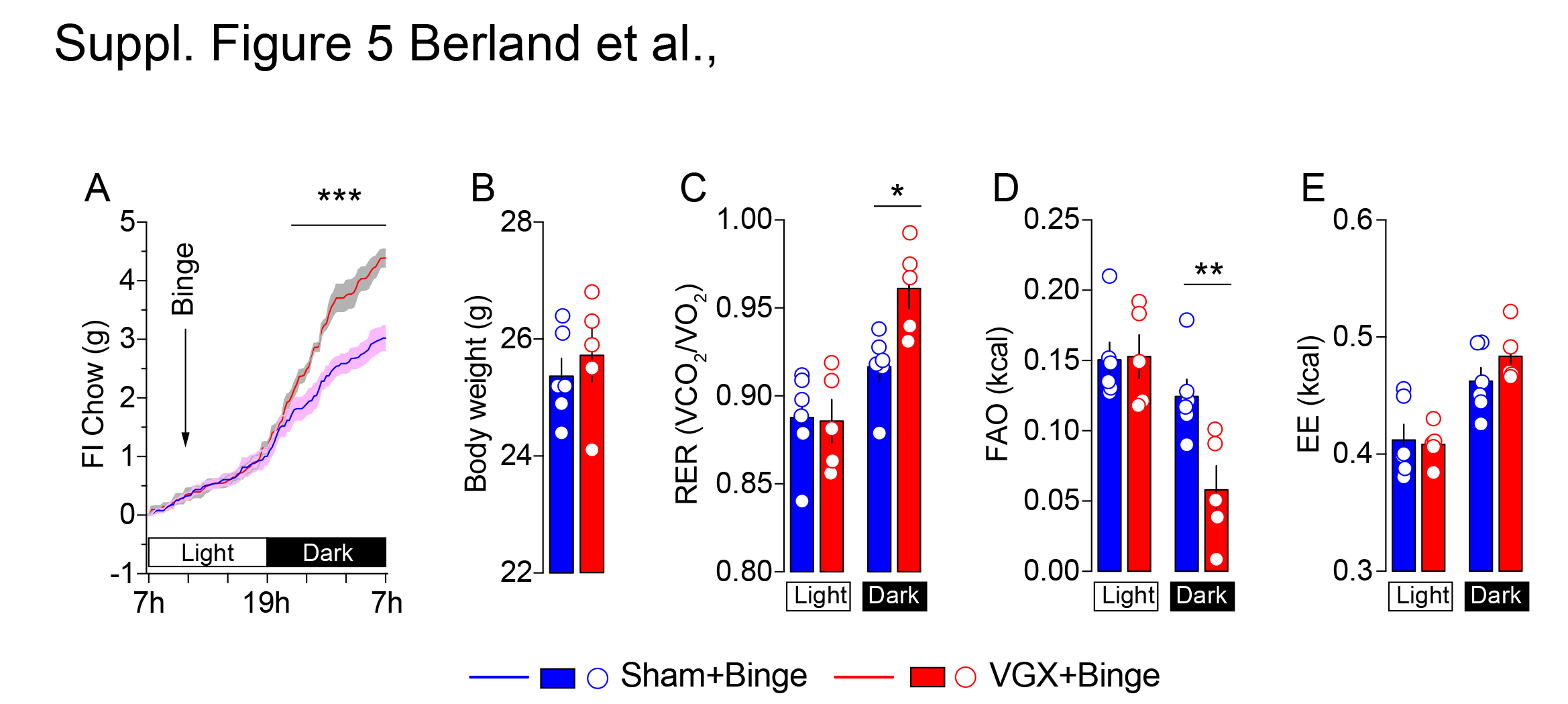

### Suppl. Figure 6

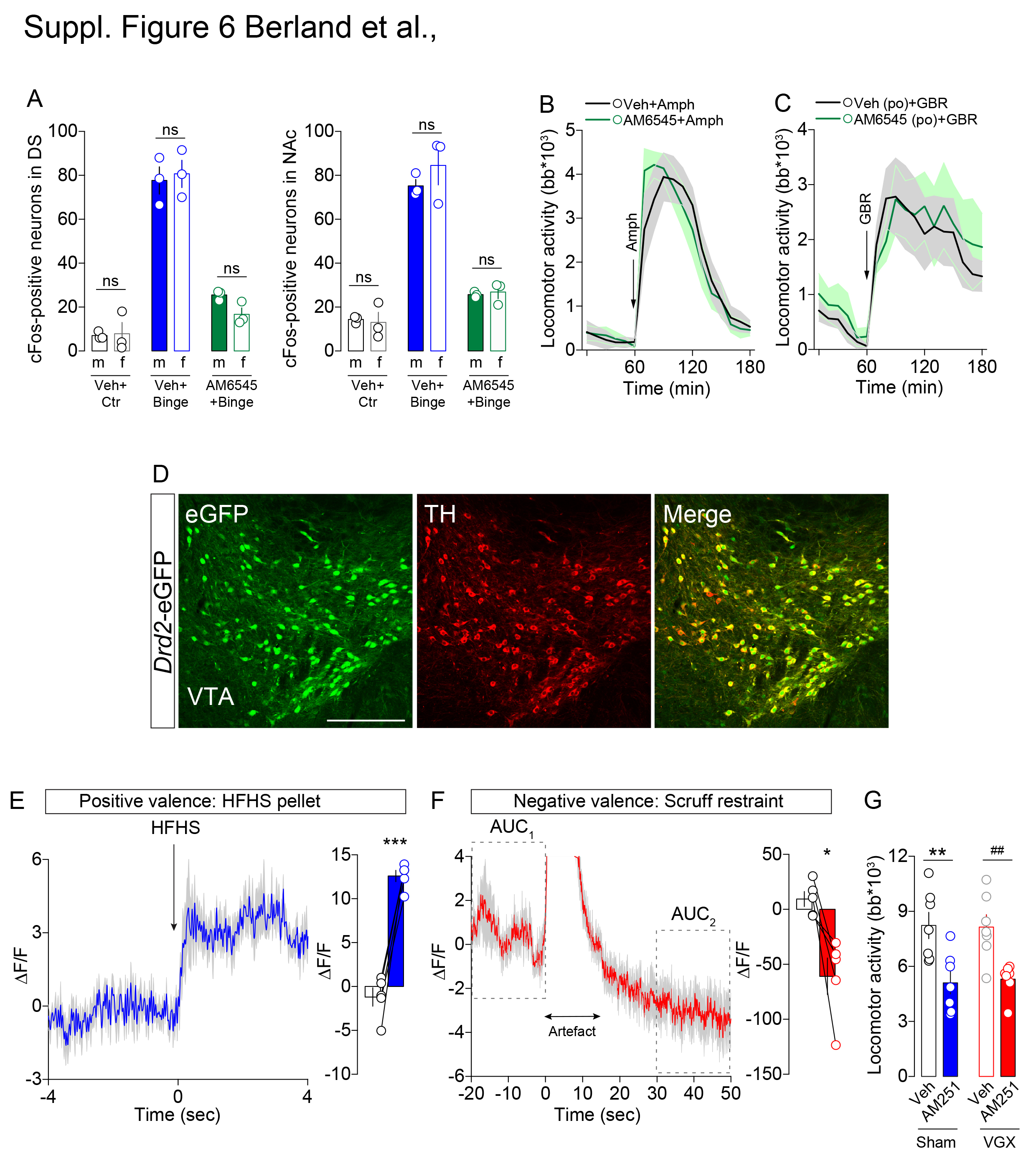
